# Systemic Factors Affect Bone Health in SMA Type II Patients and a Mouse Model of SMA

**DOI:** 10.1101/2025.02.05.636673

**Authors:** Fiorella Carla Grandi, Sonia Pezet, Alice Arnould, Sabrina Mazzucchi, Elèna Gidaja, Stéphanie Astord, Maud Chapart, Stéphane Vasseur, Alessandra Ricupero, Matteo Ernu, Yasmim Sampaio, Mathilde Cohen-Tannoudji, Pierre Meunier, Sofia Benkhelifa-Ziyyat, Lofti Miladi, Raphael Vialle, Piera Smeriglio

## Abstract

Spinal muscular atrophy (SMA) is a rare developmental disorder affecting multiple tissues. Among the non-central nervous system tissues implicated in SMA is the skeletal system, including bone and cartilage. Low bone mineral density, increased numbers of fractures of the long bones and vertebra, hip pain, and scoliosis have been reported across the spectrum of SMA patients. While lack of ambulation likely contributes significantly to bone pathology, SMA patients have markedly lower bone density compared to other non-ambulatory patients with debilitating diseases such as Duchenne muscular dystrophy, suggesting that there is a cell-intrinsic contribution of SMN to bone homeostasis and function. Mouse models of SMA have also confirmed the presence of bone and cartilage phenotypes. These alterations frequently persist post-treatment. Recent advancements in therapeutic strategies, approved by both the FDA and the EMA, have represented a leap forward in the management of SMA. However, treatment gaps remain. Post-treatment, patients frequently face continued challenges with scoliosis, bone fractures, and persistent muscle weakness—conditions that underscore the urgent need for more comprehensive therapeutic strategies with combination therapies that can support skeletal health. To date, no molecular map exists of the changes that occur in SMA patient bone and cartilage, impeding the ability of finding targeted therapies. To address this clinical need, we profiled the transcriptome of the vertebral bone and cartilage in a cohort of 11 Type II SMA patients who were undergoing surgery for scoliosis correction and compared them to 7 idiopathic scoliosis and 2 DMD controls. Additionally, we characterized the skeletal health of a mouse model of type I SMA. We find that multisystemic factors including liver and muscle health affect the underlying SMA bone pathology. Specifically, we detect alterations in the balance between osteoclasts and osteoblasts, changes in PPAR**γ** signaling, mitochondrial oxidative phosphorylation and fatty acid beta-oxidation, and alterations in the muscle-derived factor Irisin that play a role in overall SMA bone pathology.

## INTRODUCTION

Spinal muscular atrophy (SMA) is a rare developmental disorder affecting multiple tissues. Caused by the homozygous loss or deletion of the gene *SMN1*, it has been classically thought of as a motor neuron disease (MND), but increasing evidence suggests that the *SMN* protein has a role in many, if not all, tissue in the body, and that motor neurons are merely the most affected by the disease (*1–4*). The presence of a homologous gene, *SMN2*, partially compensates for the loss of *SMN1*. *SMN2* is a nearly identical copy of *SMN1*, situated in the same 5q13 locus. However, a few bases differ between the two genes, and these modify a splice junction inside of *SMN2,* which results in the production of only 10% full-length transcripts, while the remaining 90% of transcripts contain a truncated SMN mRNA without exon 7, termed *SMN*Δ*7*. This transcript is unstable, has a shorter half-life, and cannot fully compensate for the function of *SMN1* (*1*, *5*). However, *SMN2* is present in varying copies across different genomes, and thus *SMN2* copy number is the strongest genetic modifier of SMA. The severity of symptoms and the age of onset generally scale inversely with *SMN2* copy number (*1*, *5*). This variability in *SMN2* copy number, among other factors, creates a wide clinical spectrum of SMA phenotypes (*6*), which is categorized from type I to IV based on the age of onset, locomotor function, and the number of *SMN2* copies, with type I being the most severe form which is lethal in infants without treatment (*1*).

Recent advancements in therapeutic strategies, approved by both the FDA and the EMA, have represented a leap forward in the management of SMA, particularly for Type I patients. Currently, SMA Type I patients can be treated with the gene therapy onasemnogene abeparvovec (Zolgesma; Novartis) (*7*), or one of two small-molecule based therapies: the antisense oligonucleotide (ASO) nusinersen (Spinraza; Biogen) or the small molecule risdiplam (Evrysdi; Roche). Both non-gene therapy treatments alter the splicing of the *SMN2* copies naturally found in the patients’ genome and can also be used for the other subtypes (II-IV). These three treatment innovations, centered on augmenting full-length SMN protein levels by various means, have been instrumental in achieving unprecedented progress in locomotor function and extending ventilator-free survival rates (*1*, *7*). However, despite these undeniable successes, treatment gaps remain. Treated patients, especially Type II adolescents and adults, frequently face continued challenges with scoliosis, bone fractures, and persistent muscle weakness—conditions that underscore the urgent need for more comprehensive combination therapeutic strategies that can target peripheral and non-central nervous system (CNS) tissues.

Among the non-CNS tissues implicated in SMA is the skeletal system, including bone and cartilage (*2*). Low bone mineral density(*4*, *8*, *9*), increased numbers of fractures of the long bones and vertebra (*10–15*), hip pain (*16–18*), and scoliosis(*19–23*) have been reported across the spectrum of SMA patients (Types 1 to 3). Patients with SMA also present with higher bone resorption markers (*24*, *25*). While lack of ambulation likely contributes significantly to bone pathology (*24*, *26*), SMA patients have markedly lower bone mineral density compared to other non-ambulatory patients with debilitating diseases such as Duchenne’s muscular dystrophy (DMD) (*27*), suggesting that there is a cell-intrinsic contribution of SMN to bone function. In support of a role for bone and cartilage in generalized SMA pathology, the SMA-MAP project, which looked for plasma protein biomarkers of SMA, found increased levels of the cartilage and bone proteins COMP, LUM, SPP1, and CLEC3B (*28*). Indeed, the top 12 markers associated with patient outcomes were associated with Ehlers-Danloss syndrome, a connective tissue disorder, and juvenile osteoarthritis (*28*). Furthermore, these skeletal health problems do not disappear with treatment. Recent reports have shown that SMA patients treated with either the ASO or gene therapy strategy continue to have documented bone health problems, including early onset scoliosis, kyphosis, rib deformity, pelvic obliquity, and hip subluxation (*29–31*), highlighting the need to understand why skeletal problems arise in SMA.

Mouse models of SMA have also confirmed the presence of bone and cartilage phenotypes. An acute knockdown of *Smn1* in mesenchymal stromal cells (MSCs) results in a loss of differentiation capacity towards the osteogenic lineage. Furthermore, this defect is *SMN2* copy number dependent (*32*). In the Taiwanese model (FVB.Cg-Tg (SMN2)2Hung Smn1*^tm1Hung^*/J), postnatal day 1 (P1) pups that are pre-symptomatic for the neuromuscular defects were shown to already have impaired bone growth and low bone mineral density in the vertebrate column due to a chondroblast impairment (*33*). An increased rate of osteoclastogenesis and decreased osteoblast differentiation was found in SMA mice from the same model (*34*, *35*). Furthermore, the *SMN*Δ*7* isoform of *SMN* has been shown to interact with osteoclast stimulating factor (OSF) and promote formation of osteoclasts in culture, which would increase the turnover of bone (*36*). Thus, the findings from these mouse models support clinical data about the ubiquity and early onset of skeletal health problems in SMA and has paved the way for our appreciation of the role of bone health in the complex slate of SMA pathology.

However, to date, no molecular map exists of the changes that occur in the bone or cartilage tissue of SMA patients, especially how skeletal health evolves after treatment with the current state-of-the-art therapies. This lack of molecular targets prevents researchers and clinicians from making informed choices about how to develop combination therapy strategies that can address skeletal health needs, a key priority of the SMA community addresses in the 2024 SMA Europe research targets (*37*). To address this critical need, we profiled the transcriptome of the vertebral bone and cartilage in a cohort of 11 SMA Type II adolescent patients who were treated with Nusinersen. These patients were undergoing surgery for scoliosis correction; thus, the cohort reflects patients with known bone involvement. We compared these Type II patients with other adolescents diagnosed with DMD, as undergoing surgery, or with adolescent idiopathic scoliosis (AIS) to identify bone health factors particular to the SMA pathology. Finally, we compared our findings in human tissues with a mouse model of type I SMA and observed the alteration of similar pathways. Additionally, the mouse model allowed us to investigate the contribution of other organ systems such as the liver and the muscle, in SMA bone pathology. Specifically, we find alterations in the balance between osteoclasts and osteoblasts, changes in PPARγ signaling and mitochondrial health, and alterations in the muscle-derived factor irisin playing a role in overall SMA bone pathology.

## RESULTS

### Transcriptional characterization of Type II SMA vertebral bone shows changes in intracellular transport, cytoskeletal organization, and RNA processes compared to AIS and DMD controls

Despite the characterized involvement of the skeletal system in SMA, to date, there are no molecular maps of the changes that occur in human SMA bone and cartilage, impeding targeted treatment strategies. To overcome this barrier, we collected bone and cartilage from a cohort of 11 SMA Type II patients treated with nusinersen (**Figure 1A-C**). These samples were recovered from the medical discard generated during procedures to surgically correct scoliosis via a minimally invasive fusionless surgery (*38–40*) (**Supplemental Figure 1A**). In addition, as controls, we obtained the bone from 7 non-NMD controls who were also undergoing scoliosis surgery (*41*) and had a diagnosis of adolescent idiopathic scoliosis (AIS) and two patients with an unrelated neuromuscular disorder, Duchenne’s muscular dystrophy (DMD), which also has document skeletal health challenges (*42*). We reasoned that since these DMD patients have a non-SMN related muscle-disease and loss of ambulation, they would allow us to control for locomotion specific effects, if any. From a subset of these samples, we were able to also collect cartilage tissue (n=7 AIS controls, n=2 DMD, n=4 SMA). Samples were sex and aged matched to the degree possible (**Figure 1B**), however, SMA patients were on average 2-years younger than the other two groups (**Figure 1C**).

**Figure 1:**
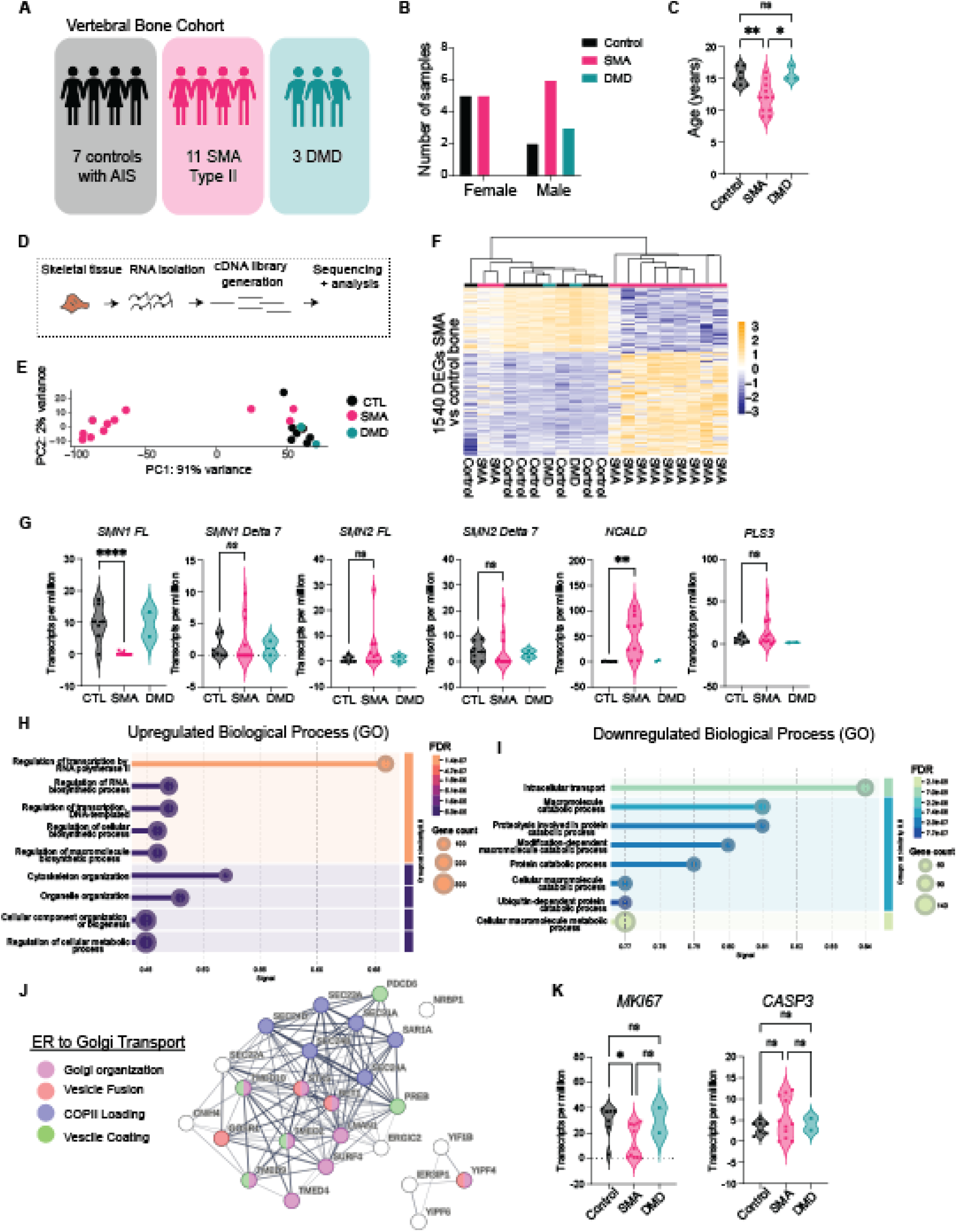
Transcriptomic Characterization of Type II SMA Vertebral Bone. **A.** Diagram of the vertebral bone cohort collected of 7 controls with adolescent idiopathic scoliosis (AIS), Type II spinal muscular atrophy (SMA) or Duchenne’s Muscular Dystrophy (DMD). Of the 3 DMD patients, only two were sequenced. **B.** Sex breakdown for each category. **C.** Age, in years, for each cohort. Means are tested via a one-way ANOVA and multiple-comparisons testing. * <0.01, **<0.001, n.s – not significant. **D.** Schematic of the RNA-sequencing workflow for the samples. **E.** Principal component analysis (PCA) of the transcriptome of each sample, which is represented by a point. The PCs were calculated using the top 2000 most variable genes. PC1 explains 91% of the variance and PC2 2% of the variance. **F.** Heatmap of the 1540 differentially expressed genes between the AIS controls and the SMA samples. Unsupervised hierarchical clustering is shown at the top. The gene expression is z-scored across each row. **G.** Top internal cell structures associated with the upregulated and downregulated gene. **H.** Violin plots shown the transcripts per million (TPM) for SMN1 full-length (FL), SMN2 FL, SMN1 delta7, SMN2 delta 7 and the two known SMA modifier genes NCALD and PLS3. Means are tested only between control and SMA samples using students t-test. * <0.01, **<0.001, n.s – not significant. DMD samples are plot as reference but were not statistically evaluated. **I.** Top biological Gene Ontology (GO) processes associated with the genes upregulated in SMA samples. The number of genes in each term is given by the size of the bubble eat the end of each bar, while the shading od each term gives the false discovery rate (FDR) and the strength of the enrichment is plotted along the x-axis. **J.** Top biological Gene Ontology (GO) processes associated with the genes downregulated in SMA samples. **K.** STRING network analysis of the genes associated with the ER to Golgi transport GO term. Each ball represents a single gene that is downregulated in SMA samples and is colored by its associated process. Nodes are connected by the known interactions, either physical or in the same network, that are known, and the thickness of the lines delineates the confidence of this interaction.

RNA was extracted from the tissue samples and utilized for generating RNA-sequencing libraries (**Figure 1D**). Our dissection of cartilage and bone from these surgical discards yielded samples with tissue-specific gene expression patterns. We compared control cartilage and bone tissues against each and found 3043 differentially expressed genes (DEGs) between the two tissues (**Supplemental Figure 1B-C**). Pathway analysis on the gene upregulated in the bone samples revealed an enrichment in hematopoietic processes (**Supplemental Figure 1D**) suggesting that the vertebral bone derived transcriptomes contain the combination of both bone-marrow and its hematopoietic populations and mesenchymal populations. However, despite this, we could observe the overexpression of classic chondrocyte markers *Col2a1* and *Sox9* in the cartilage tissue, and the osteoblast regulator *Runx2* in the bone tissue (**Supplemental Figure 1E**). The level of *Col1a1* was not different between the two tissue transcriptomes (**Supplemental Figure 1E**). Suspecting that our vertebral bone transcriptome contained a mixture of cell types, we next used a previously published single-cell RNA-seq (scRNA-seq) map of the human bone marrow (*43*) to deconvolute our transcriptome data (*44*, *45*). In addition to hematopoietic and endothelial cells, this scRNA-seq map also contained mesenchymal lineage cells (**Supplemental Figure 1F**). We observed that overall, the profile of our transcriptome was estimated to be about 25% mesenchymal, 35% endothelial and 50% hematopoietic (**Supplemental Figure 1G**), although this likely underestimates the contribution of mature osteoblasts and osteocytes that were not found in the scRNA-seq map. The proportion of mesenchymal cells was not significantly different between AIS, DMD and SMA samples, however, SMA Type II samples had a lower estimated hematopoietic fraction, with an increased endothelial fraction (**Supplemental Figure 1H**).

Principal component analysis (PCA) of the SMA, DMD and control sample transcriptomes showed that the vast majority (8/11) of SMA samples had a transcriptome highly different from the DMD and SMA samples, as also evidenced by the hierarchical clustering performed on the differentially expressed genes (**Figure 1E-F**). No metadata which we had access to were associated with the three samples whose transcriptional profiles resemble the controls more than the SMA patients. We performed differential expression analysis between SMA samples and controls and observed 938 up and 602 downregulated genes (**Figure 1F**). We also tested for a sex-effect, by looking for differentially expressed genes between the male and female SMA bone samples in our cohort but had not differentially expressed genes. However, it is important to note that the size of this cohort is not large enough to detect subtle sex-effects.

To understand the expression of SMN in bone tissue, we first turned our attention to the SMN levels. Our previous report on Type II SMA muscle showed that in Type II adolescent individuals, there is no difference in the amount of SMN full-length (SMN FL) expressed between SMA and control samples, both at the RNA and protein levels (*46*). We observed the expected lack of reads mapping to the *SMN1* locus (**Figure 1G**), however, we did not observe the same level of compensatory SMN2 FL reads (**Figure 1G**). Various attempts to perform a western blot for SMN from the remaining bone tissue were not successful, likely due to the relative low expression levels of SMN relative to other predominant proteins in the bone such as collagen. As previously noted for the muscle, there was an increase in *NCALD* expression in SMA samples but no changes in the *PLS3* levels (**Figure 1G**).

Pathway analysis (**Figure 1 H-I**) showed changes in RNA processing and cytoskeletal pathways among the upregulated genes, while changes in catabolic and endosomal membrane pathways were enriched among the downregulated genes. SMN has been previously found to interact with caveolin-1 (*47*) and alpha-COP (*48*, *49*), suggesting the reasons for the alterations win the endosomal membrane pathways, and the changes in the ER to Golgi transport (**Figure 1J**). As expected, due to SMN’s previous links to cell cycle progression, we observed a slight decrease in the amount of *MKI67*, but no difference in the cell death marker *CASP3* (**Figure 1K**). Overall, transcriptional analysis showed a vastly transformed gene regulation network, with few similarities to the DMD samples, supporting the hypothesis of SMN-dependent alterations beyond just those found in loss-of-ambulation.

### Few transcriptional changes are found in SMA Type II vertebral cartilage compared to AIS and DMD controls

We next turned our attention to the cartilage samples collected (n=7 AIS, n=4 SMA, n=2 DMD), performing the same analysis scheme as for the vertebral bone (**Figure 2A**). The vertebral bone develops via endochondral ossification and these growth plates continue to elongate during the growth spurts associated with adolescence(*50*, *51*). Thus, we reasoned that if the changes we observed in the bone tissue were due to changes occurring in the growth plate, we might be able to observe some of the alterations already occurring in the cartilage. Unlike in the bone data from above, PCA analysis did not reveal clear differences between the AIS/DMD transcriptomes and the SMA samples (**Figure 2B**). We observe the expected lack of reads mapping to the *SMN1* locus and no compensatory increase in full-length or delta7 reads from the *SMN2* locus (**Figure 2C**). Additionally, unlike in the bone, we did not observe an increase in *NCALD* expression and could not reliably detect the expression of *PLS3* (**Figure 2C**). Differential expression analysis revealed 118 DEGs, with the majority of these related to overexpression of mitochondrial DNA genes in 4 SMA samples (**Figure 2D, Supplemental Table 1**). Among these 118 DEGs, there was little overlap with the DEGs in the bone, with only 6 shared targets (**Figure 2E**). Among the differentially expressed genes, we did observe the downregulation of 5 pro-chondrogenesis genes: *COLGALT2, ANGPTL2, NKX3.2 TENT5A* and *CLIP2* and the upregulation of *DOT1L*, which has previously been associated with a protective function in cartilage physiology (*52*) (**Figure 2F**). Due to the limited size of the cartilage cohort, we could not perform sex analysis, as the cohort is not large enough to observe any differences. Overall, these results lead us to conclude that within our cohort the vertebral SMA Type II bone and bone marrow environment was more severely affected than the vertebral cartilage.

**Figure 2:**
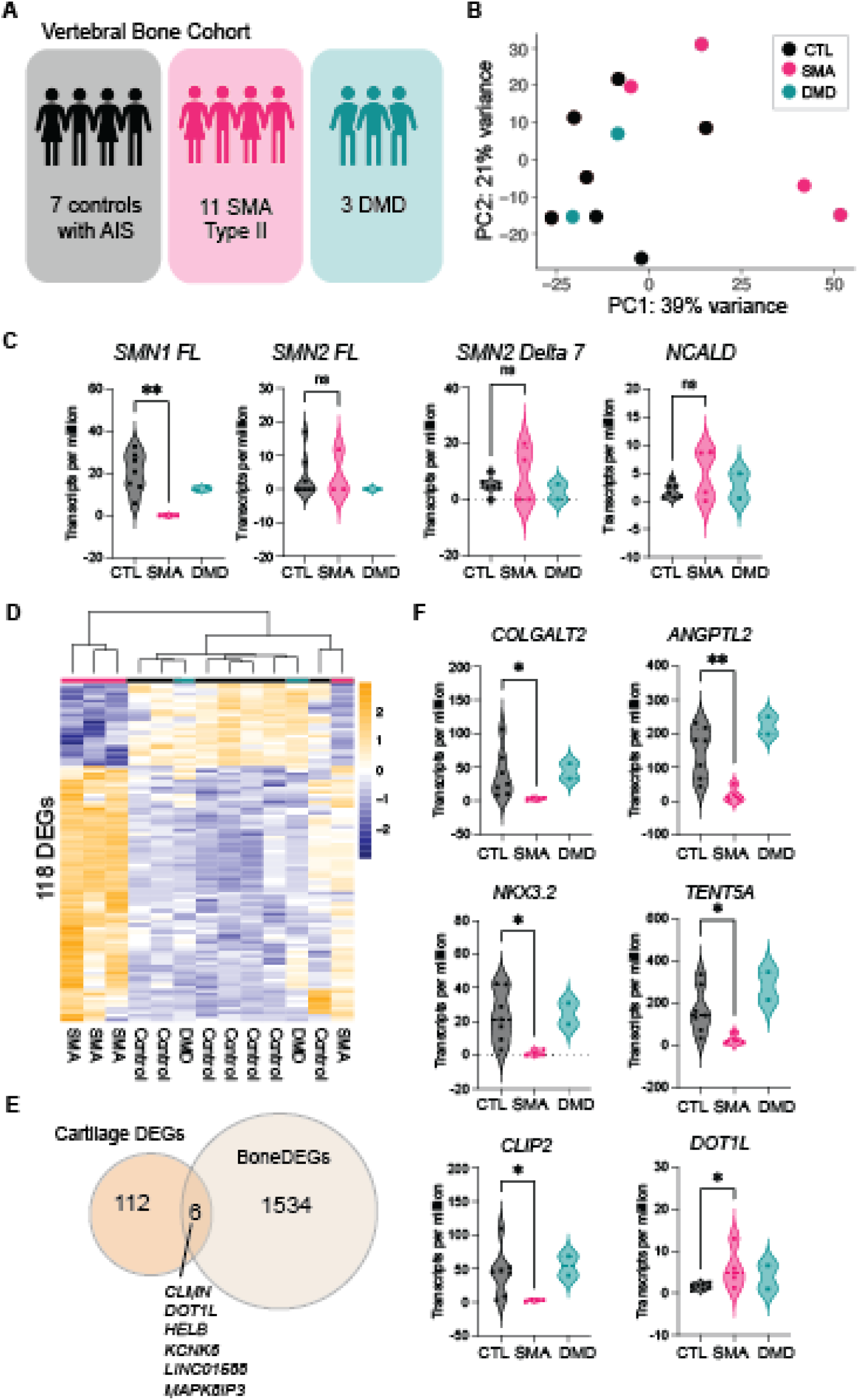
Transcriptomic Characterization of Type II SMA Vertebral Bone. **A.** Diagram of the vertebral bone cohort collected of 7 controls with adolescent idiopathic scoliosis (AIS), Type II spinal muscular atrophy (SMA) or Duchene’s Muscular Dystrophy (DMD). All cartilage samples matched to a sample from Figure 1A, however, we were not able to get both bone and cartilage from all samples. **B.** Principal component analysis (PCA) of the transcriptome of each sample, which is represented by a point. The PCs were calculated using the top 2000 most variable genes. PC1 explains 39% of the variance and PC2 21% of the variance. **C.** Violin plots of the transcripts per million (TPM) for SMN1 full-length (FL), SMN2 FL, SMN2 delta 7 and the known SMA modifier genes NCALD. *SMN1* Delta 7 and *PLS3* were not well detected. Averages are tested only between control and SMA samples using students t-test. **<0.001, n.s – not significant. DMD samples are plot as reference but were not statistically evaluated. **D.** Heatmap of the 118 differentially expressed genes (DEGs) between the AIS controls and the SMA samples. Unsupervised hierarchical clustering is shown at the top. The gene expression is z-scored across each row. **E.** Venn diagram of the DEGs that overlap between the SMA bone and cartilage. Genes with the same directional change are noted. **F** Violin plots of the transcripts per million (TPM) of the pro-chondrogenesis genes *COLGALT2, ANGPTL2, NKX3.2, TENT5A, CLIP2* and *DOT1L.* Means are tested only between control and SMA samples using students t-test. * <0.01, **<0.001, n.s – not significant.

### The SMNΔ7 mouse model shows alterations in endochondral ossification processes after symptom onset postnatally but not during embryonic development

Human samples, while powerful in detecting real-world changes associated with the disease, have limitations due to sampling size and variability, and the inability to assess the tissues longitudinally. Therefore, to get a better molecular assessment of the changes occurring in the skeletal tissues over time, we turned to a mouse model of SMA, the *SMNΔ7* strain (*53*, *54*). Bone can be formed through two distinct processes, intramembranous ossification which does not require a cartilage template, or endochondral ossification, which does. To assess which of these processes was affected, and when, we performed whole-mount skeletal staining at three time points during disease progression: post-natal day three, before obvious motor deficits, at P7, the onset of motor deficits but before the onset of extreme weight-loss and at P14 the end-stage of the disease when animals are paralyzed (**Figure 3A**).

**Figure 3:**
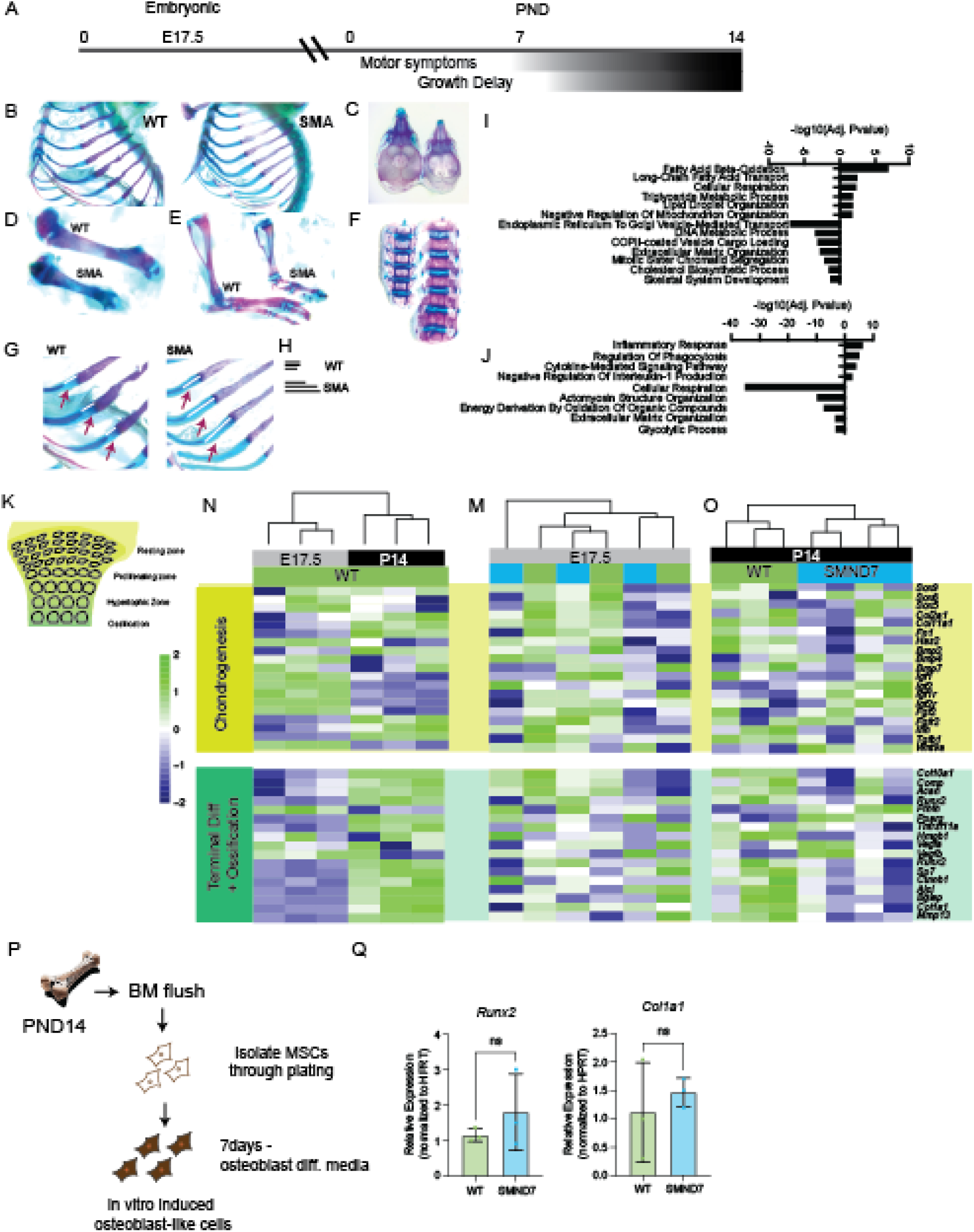
Alteration in growth plate dynamics in a mouse model of Type I SMA. **A.** Timeline schematic of the onset of symptoms in the *SMNΔ7* model. **B-H.** Whole skeletal staining of WT and *SMNΔ7* mice at postnatal day 14. In H, the length of the distance between the ossifying rib bones in (G) are shown. The smaller the size of the bar the closer the two growth plates are to each other, which is what would be expected in normal endochondral development. **I.** Pathway analysis for the genes differentially expressed in the growth plate. **J.** Pathway analysis for the genes differentially expressed in the bone transcriptome. **K.** Schematic of the growth plate in the mouse, showing the different stages of endochondral ossification. In general, the growth plate is the most immature on the edges and most mature, and ossified, in the center. **N.** Heatmap of selected genes involved in chondrogenesis and terminal differentiation and ossification at the growth plate at E17.4 and P14 in WT mice. **M.** Heatmap with the same gene set as in O but plotted for WT or *SMNΔ7* mice at E17.5. Unsupervised hierarchical clustering of the expression profiles shows that there is not clustering via genotype. **O.** Heatmap with the same gene as in O but plot for WT or *SMNΔ7* mice at P14. Unsupervised hierarchical clustering of the expression profiles shows clustering by genotype. In all heatmaps, the gene expression is z-scored across each row. **P.** Schematic of the process of isolating and differentiating mesenchymal stromal cells (MSCs) and osteoblastic differentiation. **Q.** qPCR for the osteoblastic genes *Runx2* and *Col1a1* after MSC differentiation in WT and SMA animals. Means of each genotype were tested via a student’s t-test. N.s.- not significant.

Skeletal staining at P3 and P7 showed no obvious deficits in the skeletal development, both among the endochondral and intramembranous bones of the face and skull (**Supplemental Figure 2A**). By contrast, by P14 there were clear changes in the skeletal morphology, with animals having much smaller bones and ribs (**Figure 3B-H**), consistent with the overall decrease in body weight. In addition, the delay in endochondral ossification was particularly clear in the growth plates of the ribs, where the ossified region was smaller in SMNΔ7 animals than in littermate controls (**Figure 3G-H**). Bones formed through intramembranous ossification, like the calvaria, appeared to be of normal morphology, by contrast, suggesting an exclusive problem with the endochondral ossification process in the postnatal animals (**Figure 3E**). A recent report using a Prx1-Cre mouse showed that selective loss of *Smn1* in mice causes the formation of small bones, and that this phenotype can be rescued in a dose dependent manner by *SMN2* (*32*). To rule out any early developmental delays in endochondral ossification, we dissected the growth plate cartilage and the bone of mice at embryonic day 17.5 (E17.5), when the growth plate is forming, and the hypertrophic area is being established. Immunofluorescence of COLX showed no differences in the development of the growth plate (**Supplemental Figure 2B**). Additionally, several classic molecular markers of the endochondral process were unaltered (**Supplemental Figure 2C).**

To get a better assessment of the molecular pathways altered in the processes of endochondral ossification and to compare them to those found in our patient cohort, we performed RNA-seq on the growth plate cartilage and mature bone, including all the zones of the development growth plate at E17.5 and P14. As expected, we observed the total absence of transcripts derived from the mouse *Smn1* in SMA animals (**Supplemental Figure 2D**). PCA analysis of the transcriptomes showed vast differences between the E17.5 and P14 growth plate and bone, and PC2 captured the effect of genotype at P14 (**Supplemental Figure 2E**). No differentially expressed genes were found between SMNΔ7 and WT littermates at E17.5, while 1948 DEGs were observed between SMNΔ7 and WT littermates in the growth plate and 632 DEGs were found in the P14 bone (**Supplemental Figure 2F, G).** Among the upregulated pathways in the growth plate, we found terms related to fatty acid oxidation, cholesterol processing, and metabolism (**Figure 3I**). Among the downregulated genes in the growth plate, we found processes related to cell division and mitosis, DNA replication and the endoplasmic reticulum to Golgi vesicle-mediated transport. Specifically, we also observed the downregulation of *PCNA, Ccne2, Cks1b, Ccna2, Ccnb1,* and *Cks2* which are associated with the procession of cells between the G1 and S phases of the cell cycle (**Supplemental Figure 2H)**. This effect is consistent with the known roles of SMN in the cell cycle. This alteration in the cell cycle likely leads to difficulties with growth plate condensation, as chondrocytes needs to expand massively. The other notable pathway that was downregulated was related to the endomembrane system (**Figure 3I**), as we had also observed in the DEGs of the SMa bone tissue. Systematic analysis of the genes associated showed a decrease in the COPI mediated transport between the Golgi and ER as all as changes in the COPII system (**Supplemental Figure 2I)**. SMN has been previously shown to directly interact with COPI member α-COP, suggesting that the absence of this interaction may alter the endomembranous system in the growth plate(*48*, *49*). We observed that α-COP (*Copa)* was downregulated at P14 (**Supplemental Figure 2J)**. This alteration of the ER and Golgi transport processes may alter the secretion of the various collagens, which are dependent on these processes for secretion.

The growth plate cartilage in the SMA animals represents a way to understand how endochondral ossification is altered – something we were not able to assess from our human samples. Therefore, we focalized our analysis on a series of genes known to control the process of endochondral ossification, which occurs through a delicately orchestrated processes with various stages (**Figure 3K**). As expected, between WT animals at E17.5 and P14 we saw a switch between the expression of predominantly chondrogenic genes at E17.5 when the cartilage template is being established and a higher expression of terminal differentiation and ossification genes at P14 when the cartilage template is giving way to bone (**Figure 3N**). We next compared these gene sets at E17.5, and as expected, observed no clear clustering of samples by genotype (**Figure 3M**). By contrast, at P14 we could observe a clear clustering of samples by genotype, and a decrease in both chondrogenic genes but predominantly a decrease in terminal differentiation and osteogenesis genes, among these the down regulation of *Col1a2, Col10a1 and Mmp13* (**Figure 3O**). Conversely, we observed the upregulation of *Pparg* and *Pthlh*, whose upregulation is associated with a delay in endochondral development (*55*, *56*) (**Figure 3O**).

We were surprised to note that the terms related to the bone differentially expressed genes were not necessarily related to skeletal development, as in the growth plate, but on cytokine-related signaling and inflammation, among the upregulated, and oxidative phosphorylation and glycolytic process, among the downregulated. (**Figure 3 J**). This, in combination with the alterations of the growth plate which we only observed at the late time point, led us to hypothesize that prior to the switch in whole body growth that appears between P7 and P14, the bone was developing normally, and that at P14 we are now observing reactions to systemic alterations. Few genes were commonly differentially expressed between the two tissues, further underscoring the different reaction of cartilage and bone to the chronic reduction in SMN expression (**Supplemental Figure 2K**). To further validate this hypothesizes that osteogenic lineage commitment is not impaired in the SMNΔ7 mice, we isolated mesenchymal stromal cells (MSCs) from the bone marrow of P14 WT and SMA mice and differentiated these cells towards osteoblast-like cells *in vitro* (**Figure 3P**). In line with our other data, these MSC-derived osteoblast-like cells expressed equivalent levels of *Runx2* and *Col1a1* between SMA and WT mice, suggesting that osteogenic induction can occur equivalently well (**Figure 3Q**). All together, we hypothesized that the cartilage and bone begin to respond to systemic pathogenic cues in the SMA mouse sometime after P7. Given the large number of terms associated with lipid oxidation, cholesterol metabolism, mitochondrial function and the upregulation of *Pparg* in the growth plate, we hypothesized that this might be related to the known metabolic defects in the SMA models.

### ICV injected AAV9 mediated gene therapy at P1 restores bone transcriptome in the short and long-term

Recent reports have shown that even after treatment, Type I and Type II patients continue to face skeletal health challenges (*29–31*). However, these therapies are often difficult to give before symptom onset or may not target all tissues. Thus, we next wanted to determine if our previously established gene therapy approach (*57*, *58*), which can effectively expand the lifespan of the SMNΔ7 model, could restore the altered transcriptional pathways in the bone. As previously established, SMNΔ7 mice were injected ICV at postnatal day 1 with 5 × 10^10^ vg (**Figure 4A**). Our gene therapy approach simulates an idealize treatment scenario as animals are treated at P1 before symptom onset. Animals were then assessed at 7 and 14 days after gene therapy, as well as 6 months after gene therapy for a long-term time point (**Figure 4A**). Our gene therapy can effectively deliver the optimized *Smn* to key central nervous tissues (brain and spinal cord) as well as key peripheral tissues (skeletal muscle, heart and liver) by P7 (**Figure 4B**). This viral copy number decreases significantly in most tissues between post-natal day 7 and 6-months (**Figure 4B**) after injection, as is expected because the AAV genome is non-replicating and non-integrative. By P7, all tissues have some level of SMN expression, with some tissues have supraphysiological doses (brain, liver and heart) and some tissues having lower doses that WT animals, such as muscle (**Figure 4C**). At LT, these levels generally decreased from their supraphysiological levels (**Figure 4 D, E**). Thus, our AAV-therapy provides both CNS and systemic rescue to the SMA mice.

**Figure 4:**
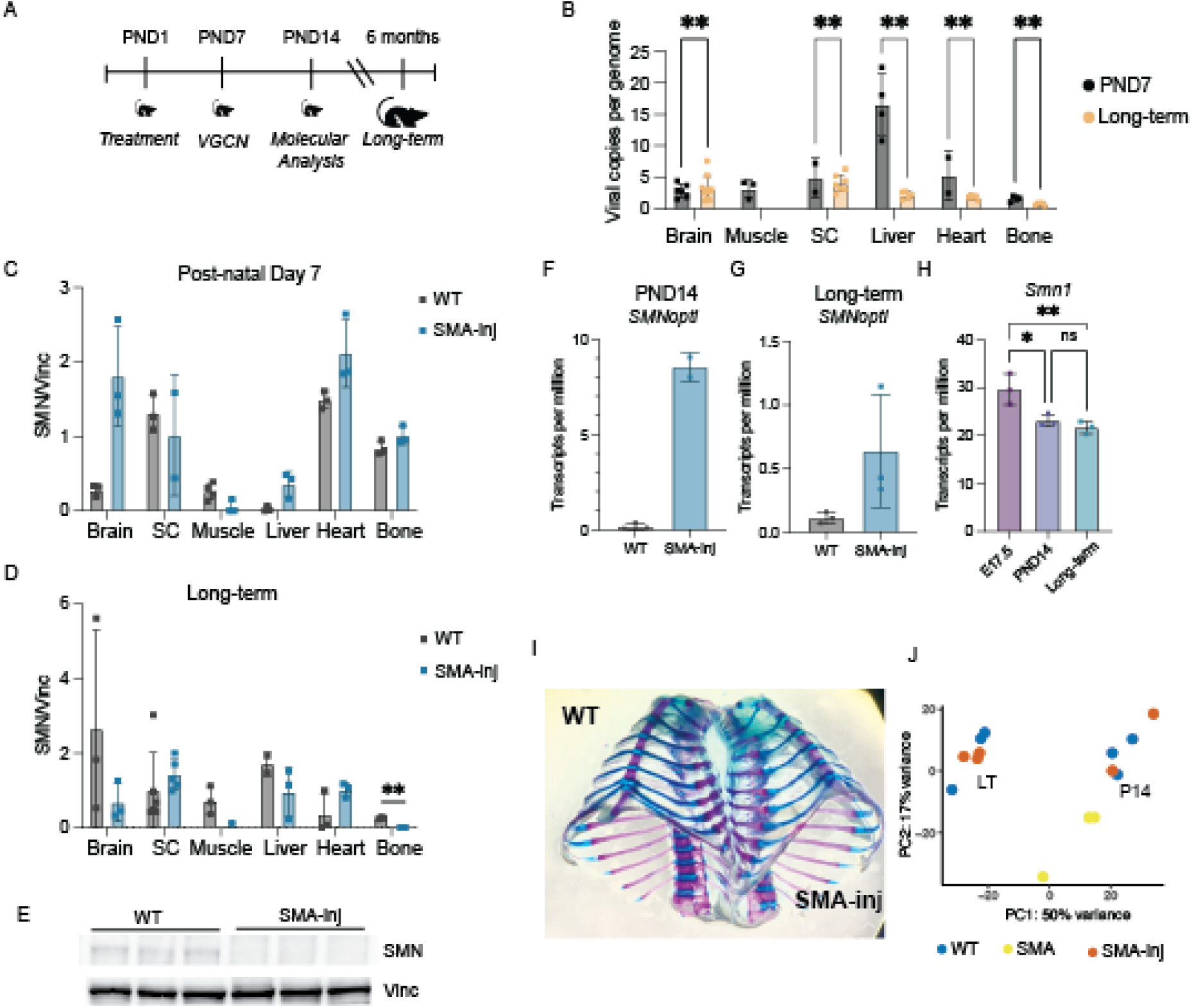
AAV9-SMN1 mediated gene therapy at P1 restores bone health in the short and long term. A. Schematic of the treatment in AAV9-SMN1 gene therapy and the different time points of collection. **B.** Viral copies per genome number (VCGN) for brain, muscle, spinal cord, liver, heart and bone tissue at P7 after injection and 6 months after injection (LT). The means of the VCGN measurements were tested via a one-way ANOVA followed by multi-comparisons testing with pre-selected pairs. P7 and LT measurements were compared only within a particular tissue and not between each other. ***<0.001. **C.** SMN protein normalized to vinculin levels in the brain, spinal cord, muscle, liver, heart and bone at post-natal day 7 after injection for SMA mice and in non-injected WT controls of the same age. The means of the protein levels measurements were tested via a one-way ANOVA followed by multi-comparisons testing with pre-selected pairs. WT and SMA-injected measurements were compared only within a particular tissue and not between each other. No comparisons were found to be statistically significant. **D.** SMN protein normalized to vinculin levels in the brain, spinal cord, muscle, liver, heart and bone 6 months after injection for SMA mice and in non-injected WT controls of the same age. The means of the protein levels measurements were tested via a one-way ANOVA followed by multi-comparisons testing with pre-selected pairs. WT and SMA-injected measurements were compared only within a particular tissue and not between each other. **<0.01. **E.** Representative western blot of SMN and vinculin levels in the long-term bone. **I.** Whole-skeletal staining of P14 SMA mice injected with the AAV9-SMN therapy. **J.** PCA plot of the transcriptomes of the bone tissue of P14 and long-term (6 month) injected SMA mice. No differentially expressed genes were found between WT animals and SMA injected animals. Each point represents a single animal’s transcriptome.

To determine if the skeletal system was well target by our therapy, we measured the VGCN in bone at P7 where we observed mean VGCN of 1.6 +/- 0.4. This decreased to 0.7 +/- 0.3 at 6-months. This low-level VGCN was in line with the AAV-derived transcripts we could map upon sequencing total P14 and LT bone mRNA after AAV injection, with an average of 8.5 transcripts per million at day 14 and a mean of 0.6 transcripts per million at 6-months (**Figure F, G**). Similarly, in P7 bone, we observed a restoration of the SMN protein level, even though the copies of mRNA were much lower than those found for WT mouse *Smn1 (***Figure 4H)** suggesting that there is at least some levels of translational regulation to SMN quantities. Indeed, the non-correlation between SMN RNA and protein levels has been previously described (*59*).

We next performed whole-mount skeletal staining on injected animals at P14, where we had previously observed defects in the skeletal development. Animals had wildtype-like development of the rib cage and long bones (**Figure 4I**) suggesting that gene therapy could rescue the skeletal growth defects we previously observed. To determine if the transcriptome of the bone was restored, we performed RNA-sequencing at 14-days and 6-months post gene therapy, to simulate both short-term and long-term treatment. Principal component analysis revealed that at P14, we could see a restoration of the changes observed in SMA animals at P14 (**Figure 4J**), and no differentially expressed genes were observed, except for *Smn1* and *Smn1-Opti* derived from the AAV. Similarly, in the long-term injected animals, we saw transcriptomes highly similar to the WT animals, with no differentially expressed genes aside from *Smn1* (**Figure 4J**).

We concluded that our gene therapy approach can deliver the SMN protein to the skeletal tissues during the early postnatal stages, however likely due to the low initial starting VGCN and the remodeling of the bone during normal homeostasis, these viral copies are not distributed to newly generated bone cells and protein levels are not restored in the bone in the long-term. Despite this, transcriptionally, the bone remains similar. This, in combination with the late onset of the skeletal phenotypes and our human data lead us to hypothesize that a significant component of the skeletal defects observed as due to changes in systemic factors in other tissues, which are well rescued by our therapy.

### SMA Type II vertebral bone has an enrichment for a signature of active osteoclasts

An important aspect of bone health is the balance between osteoblasts, the anabolic cells of bone tissue responsible for collagen secretion and mineralization, and osteoclasts, the catabolic cells of the tissue responsible for degrading and remodeling this matrix. Together with other resident cells, they form the basic remodeling unit (BRU) (*60*). Changes to this balance alters bone homeostasis, with increase osteoclast activity causing osteoporosis and decreased osteoclast activity causing osteopetrosis. To better assess this balance, we calculated an osteoclast and osteoblast score for each patient from our vertebral bone transcriptomic data using a previously established gene list (*61*) (**Figure 5A**). After scoring, we observed that SMA patients had an increased osteoclast score, and a decreased osteoblast score compared to controls (**Figure 5B**). We also scored the DMD samples, which were in the range of the AIS controls, although no statistical testing was performed due to the low number of samples. We reasoned that the increase in the osteoclast signature might be due to an alteration in the factors related to osteoclastogenesis, which are expressed by different components of the bone and bone marrow environment (**Figure 5C**). Therefore, we checked for the expression of RANKL (*RANKL)*, which increases osteoclast formation and is the target of many anti-resorption therapies, and OPG (*TNFRSF11B*), which is an osteoclastogenesis inhibitor. However, their expression profiles were similar between AIS controls and SMA samples (**Figure 5D**). However, we observed an increased expression of pro-inflammatory cytokines and molecules, which have been associated with enhanced osteoclastogenesis, including *IL1B, ActivinA (INHBA),* and *IL34* (**Figure 5E**). While none of these molecules is enough to drive osteoclastogenesis alone, they can potentiate the activity of RANKL (*62–69*). Collectively, this suggests that SMA vertebral bone tissues may have a pro-osteoclastic environment driven by an increase in inflammatory cytokines. We assessed our SMNΔ7 for the same osteoclast – osteoblast imbalance but found no difference between PND14 SMNΔ7 and WT animals (**Supplemental Figure 3A,C**) and no change between the injected and non-injected animals at long or short-term (**Supplemental Figure 3A-C**).

**Figure 5:**
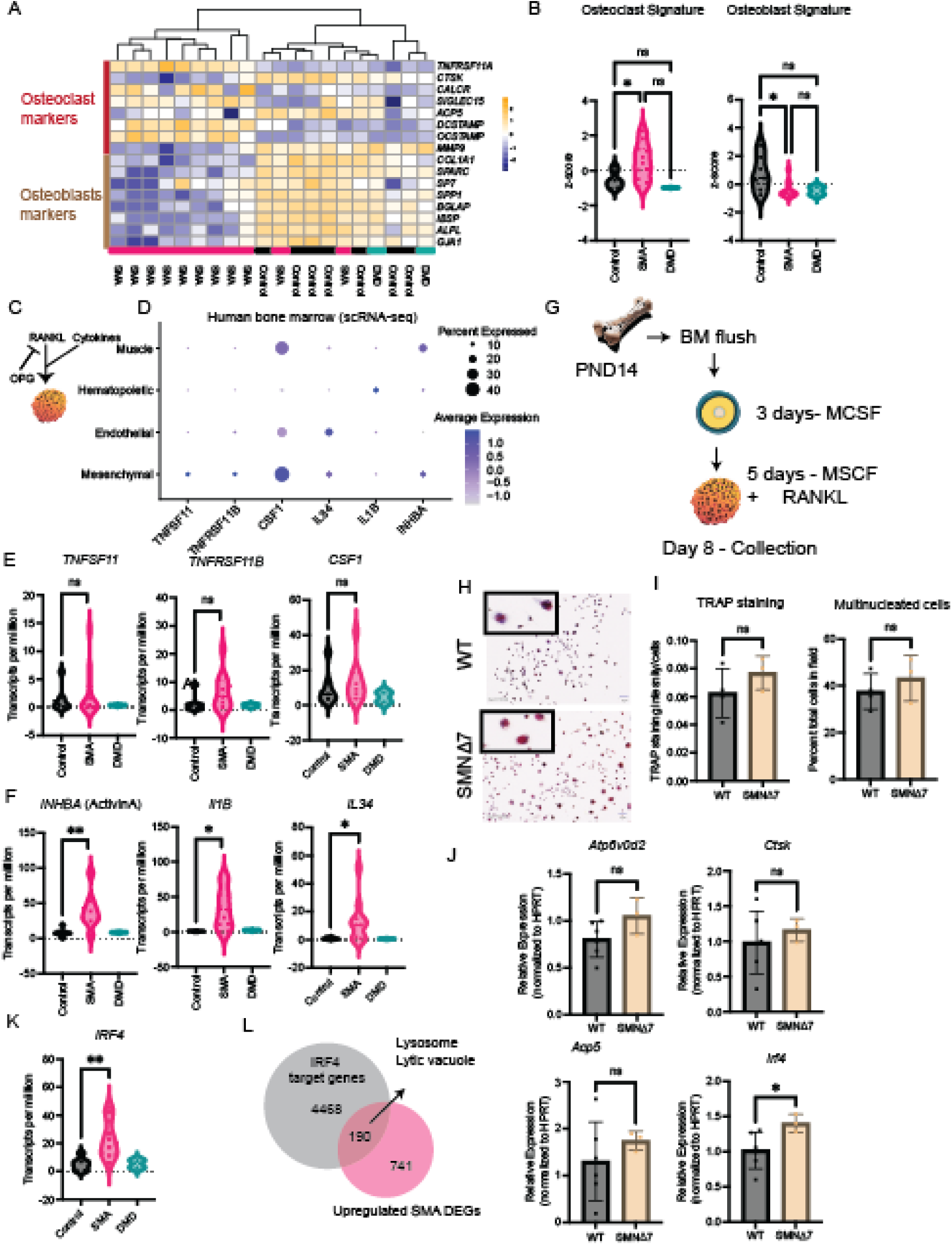
SMA Type II bone has signatured of increased osteoclast activity. **A.** Heatmap of the osteoclast and osteoblast genes used to make the respective signatures in B. **B.** Scoring of the osteoclast and osteoblast signatures in each of the SMA, AIS and DMD transcriptomes. The means of the scores were tested with a one-way ANOVA followed by multiple-comparison testing. * <0.01. **C.** Schematic of osteoclast differentiation signals, whose expression is plotted in the dot plot on the right. **D.** Human bone marrow single-cell data were reanalyzed and the expression of each osteoclast related factor is plotted for each broad category of cells. **E.** Violin plots of the transcripts per million (TPMs) of *TNFSF11, TNFRSF11B* and *CSF1.* Means are tested only between control and SMA samples using students t-test. n.s – not significant. DMD samples are plot as reference but were not statistically evaluated. **F.** Violin plots of the transcripts per million (TPMs) of *INHBA, IL1B* and *IL34.* Means are tested only between control and SMA samples using students t-test. * <0.01, **<0.001, n.s – not significant. DMD samples are plot as reference but were not statistically evaluated. **G.** Schematic of the isolation and in vitro culture of osteoclasts starting with mouse bone marrow. **H.** Representative images of the TRAP staining of the *in vitro* derived osteoclasts from WT and *SMNΔ7* mice. **I.** Quantification of the TRAP staining intensity. **I.** Quantification of the number of multinucleated cells at Day 8 of in-vitro differentiation. Means are tested via a student’s t-test *< 0.01. n.s. – not significant. **J.** qRT-PCR quantification of the expression of osteoclast marker genes at day 8 of in vitro culture. Means are tested via a student’s t-test *< 0.01. n.s. – not significant. **K.** Violin plots of the transcripts per million (TPMs) of *IRF4.* Means are tested only between control and SMA samples using students t-test. n.s – not significant. DMD samples are plot as reference but were not statistically evaluated. **L.** Ven diagram of the number of IRF4 target genes, derived from the EnrichR database, overlapping with the upregulated genes in the SMA bone transcriptome. Enriched processes in the overlapping genes are highlighted.

Previous reports have suggested that there is an increase propensity for SMA monocytes to differentiate into osteoclasts in the Smn^−/−^ SMN2 mouse model (*34*).The authors suggested this may be due to the interaction of the *SMNΔ7* isoform with osteoclast-stimulating factor(*36*). To determine if a similar propensity towards osteoclastogenesis was true in our SMNΔ7 mouse model, we isolated monocytes from P14 WT and SMA animals and cultured them *in vitro*. Cells were first pushed toward a macrophage-like state using M-CSF and then towards an osteoclast-like state using RANKL (**Figure 5F**). After 8 days in culture, cells were stained for tartrate-resistant acid phosphatase (TRAP) to determine their osteoclastic activity (**Figure 5G**). Quantification revealed a similar intensity of TRAP staining between both WT and SMNΔ7 animals (**Figure 5H**) as well as a similar percentage of multinucleated cells in each well (**Figure 5I**). To better assess the osteoclasts activity capacity, we performed gene expression analysis using real-time quantitative PCR (RT-qPCR) on the cDNA derived from the in vitro differentiated cells and observed that several osteoclast activity markers, *Cstk, Atp6v0d2* and *Acp5 (Trap)* were unchanged (**Figure 5J**). However, we observed an increase expressed in *Irf4* in SMNΔ7 osteoclast-like cells (**Figure 5J**), mirroring what was observed in the SMA vertebral bone transcriptome (**Figure 5K**). IRF4 has been previously found to potentiate osteoclast differentiation and activity(*70*, *71*). Overlapping a list of IRF4 target genes from EnrichR (*72*) with the upregulated genes in the SMA vertebral bone, we observed that 20% of the upregulated genes were IRF4 targets, pathway analysis of these genes showed an increased in terms related to lysosome function and lytic vacuoles, consistent with an increase in the phagocytotic activity of osteoclasts (**Figure 5L**). If this a direct effect of the interaction of OSF as previously proposed remains to be determined. It is important to note that the *SMNΔ7* model, even the wildtype animals have a transgenic copy of the SMNdelta7 cDNA, thus our model cannot assess the influence of increased SMND7, just the loss of the mouse *Smn1*. Altogether, our data suggests that a pro-inflammatory bone marrow environment may enhance certain cell-intrinsic aspects of SMA osteoclasts to be more active and collectively tip the balance towards a pro-osteoclastogenic signature, increasing bone weakness and porosity in Type II SMA adolescents.

### Serum from symptomatic SMNΔ7 mice turns on PPARγ signaling in human osteoblasts

We next turned our attention to common features between the human Type II vertebral bone data and the *SMNΔ7* model long-bone data, reasoning that this would allow us to study potential systematic effects **(Supplemental Figure 4A).** Overlapping both genes and pathways, we found an enrichment for processes related to fatty acid oxidation, cholesterol and fatty liver disease. In the human vertebral bone, we observed an increased expression of *CD36* and *LDLR*, two receptors that can update fatty acid and cholesterol (**Figure 6A)**, as well as an increased expression in *PPARG* and PPARA (**Figure 6A**). Indeed, overlapping the PPARG targets taken from EnrichR with the upregulated genes in the mouse or human SMA bone, we saw that 24% of the upregulated genes in human and 27% of the upregulated genes in mouse were also PPARG targets (**Figure 6C)**. Overall, these signatures matched well with our previously observed increase in *Pparg* in the growth plate, and further supported our hypothesis that the changes we observed might be due, in part, to systemic effects derived from liver metabolism.

**Figure 6:**
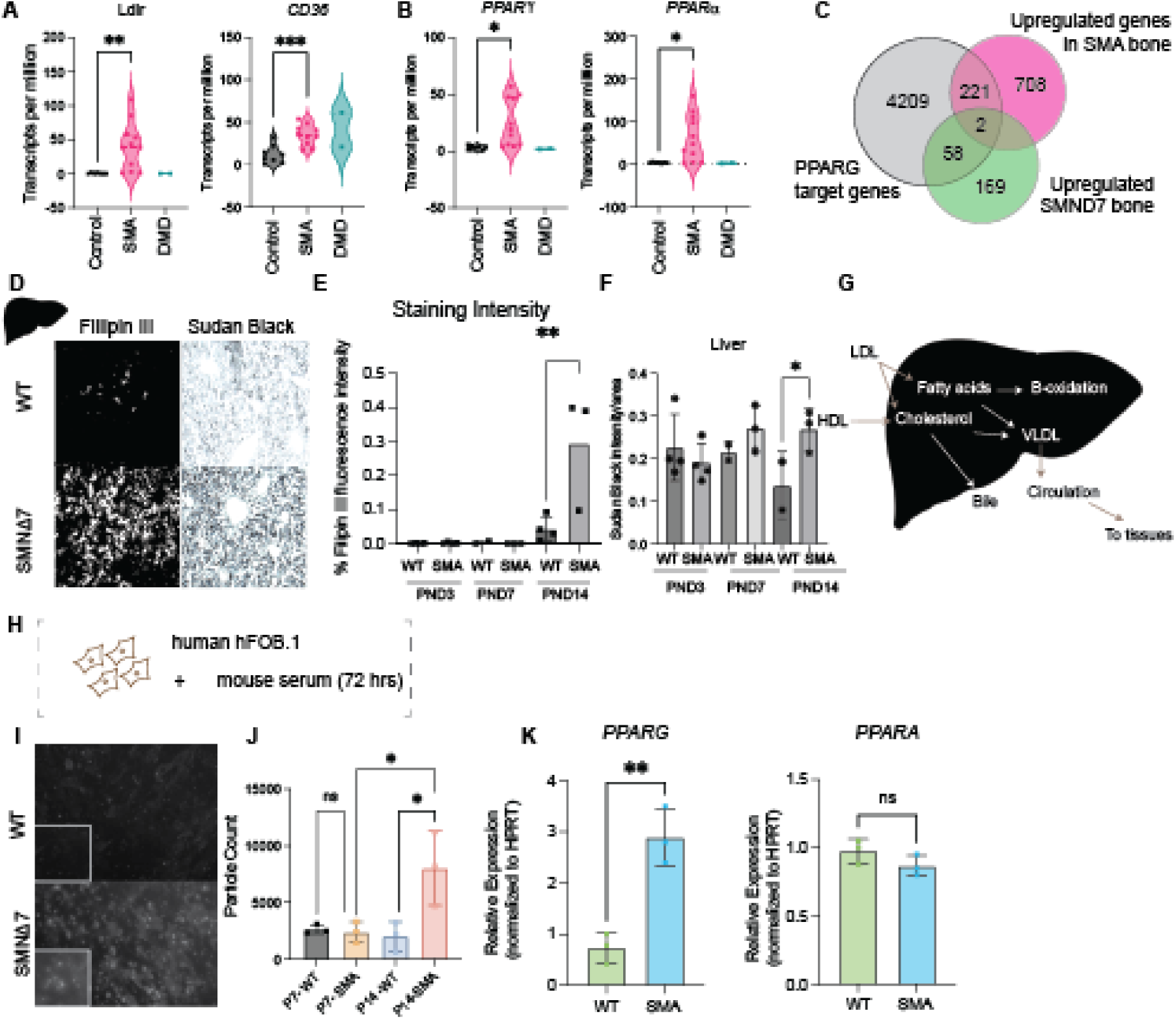
Serum of Symptomatic SMA mice causes cholesterol accumulation in PPARG activation in human osteoblasts. **A.** Violin plots of the transcripts per million (TPMs) of *Ldlr* and *CD36.* Means are tested only between control and SMA samples using students t-test. * < 0.01, ** < 0.001, ** <0.0001. DMD samples are plot as reference but were not statistically evaluated. **B**. Violin plots of the transcripts per million (TPMs) of *PPARG* and *PPARA.* Means are tested only between control and SMA samples using students t-test. * < 0.01. DMD samples are plot as reference but were not statistically evaluated. **C.** Ven diagram of the number of PPARG target genes, derived from the EnrichR database, overlapping with the upregulated genes in the SMA bone transcriptome from mouse and human samples. **D.** Fillipin III and Sudan black staining of P14 Liver from WT and *SMNΔ7* mice. **E.** Quantification of the staining intensity of Fillipin III at P3, P7 and P14. Means were compared via a one-way ANOVA with multiple comparison. **F.** Quantification of sudan black staining in the liver over time. **G.** Schematic of the processing of fatty acids and cholesterol by the liver, which are then either processed into bile acids or exit the liver as VLDL that are destined for circulation and tissue uptake. **H.** Schematic of the experimental setup treating human immortalized osteoblast line hFOB.1 with mouse serum for 72 hours. **I.** Representative image of Fillipin III staining on hFOB.1 osteoblasts treated with either P14 WT or *SMNΔ7* serum. **J.** Quantification of the staining intensity of the Fillipin III staining after osteoblast treatment with serum from P7 or P14 WT or *SMNΔ7* mice. Means were tested with a one-way ANOVA followed by multiple comparisons testing. N.s – not significant. * <0.01. **K.** qRT-PCR of the expression of PPARG and PPAR in human osteoblasts after treatment with P14 WT or SMA serum.

Lipids and cholesterol have a complex relationship with skeletal health (*73–75*). However, increasing evidence show that changes in whole-body metabolism found in obesity, type II diabetes and non-alcoholic fatty-liver disease (NAFLD/NASH) have decreased bone mineral density (*74–76*) and high levels of cholesterol have been shown to drive pro-osteoclastogenic pathways (*77*, *78*). These observations were particularly interesting to us in the context of the known changes in whole-body metabolism in SMA patients, which include increased levels of cholesterol and fatty acids in the serum and NAFLD-like symptoms in the liver of SMA models and patients (*79–85*). Therefore, we hypothesize that the liver-mediated disease, including the alteration metabolic components may be affecting the development of the cartilage and bone. To confirm the involvement of the liver in the *SMNΔ7* model, we collected livers from P3, P7 and P14 animals and stained them with Fillipin III to assess cholesterol accumulation and Sudan Black to assess fatty acid accumulation (**Figure 6D)**. Accumulation of cholesterol was seen only at P14 (**Figure 6C, Supplemental Figure 4B)**. Accumulation of cholesterol was also evident in the skeletal muscle and spinal cord using mass-spectrometry, although these levels were not high enough to be detected with Fillipin III staining (**Supplemental Figure 4D-F**). We concluded that there is an accumulation of fatty acids and cholesterol in a systemic manner in the *SMNΔ7* model, occurring between P7 and P14, between the timeframe of the motor-deficits and changes in the growth plate.

The liver is the major processing station for fatty acid and cholesterol, which are then packaged and sent into circulation where they can be taken up by different tissues for their energetic needs (**Figure 6F)**. However, an overwhelmed liver, either through excess dietary consumption or because of the liver’s inmate incapacity to process the incoming fatty acid and cholesterol, will cause increased levels of cholesterol and fatty acids in the blood, as well as increased HDL levels (*86*). This has already been observed in SMA patients and animal models (*79*, *80*). Thus, we reasoned that if the PPARγ signature that we observed in the SMA vertebral bone transcriptome was due to excess serum lipids and cholesterol, then treating human osteoblasts in culture with the serum collected from SMNΔ7 animals should increase the PPAR**γ** expression (**Figure 6H)**. Indeed, after treating human osteoblasts with P14 SMA serum we could see an increased accumulation of cholesterol using Fillipin III staining (**Figure 6I, J)**, which did not occur when cells were treated with WT serum, or serum from P7 SMA animals (**Figure 6J)**, before the accumulation was observed. We next performed qRT-PCR of the treated cells and observed an increase in PPARG but not PPARA after 72 hours to treatment, confirming our hypothesis (**Figure 6K)**.

### Mitochondrial OXPHOS changes in the bone tissue of the SMNΔ7 mouse model and Type II SMA

Another common feature between changes in the human and mouse data were pathways related to mitochondrial function. Indeed, overlapping the differentially expressed genes (DEGs) in the human bone and cartilage samples with the Human Mitocarta v3 list of genes associated with mitochondrial function (*87*), we observed that 8% of the genes were implicated (**Figure 7A)** and 10% overlap in the mouse (**Figure 7B)**. This was also in concordance with our results in Type II SMA paravertebral muscle, where we also observed a downregulation in oxidative phosphorylation(*46*). Recent work has shown that mitochondrial dysfunction can impair osteogenesis and accelerate age related bone loss (*88*). Taking the mitochondrial terms that were downregulated in the human bone samples, we observed a clustering of four predominant pathways: fatty acid beta oxidation, mitochondrial translation, particularly the ribosomal genes, cellular respiration and oxidative phosphorylation and mitochondrial organization (**Figure 7C).** Similarly, among the downregulated mitochondrial genes in the mouse bone we saw that they predominantly belong to the electron transport chain and its assembly, with most factors belong to Complex I or Complex IV (**Figure 7D).** Of these mitochondrial genes, we had an overlap of four genes, although the pathways were highly similar (*Cycs, Hadh, Timm17a and Ndufb5)*. Among the upregulated genes, we also saw some mitochondrial processes, mainly related to in the input of cholesterol or fatty acids into the mitochondria (**Figure 7E).** (**Figure 7C)** For example, the upregulation of *CPT1A* and *CPT2*, for the input of fatty acids via the carnitine cycle (**Figure 7F)** and the import of cholesterol to the mitochondria via *STAR* (**Figure 7F).** To determine if the changes in the mitochondrial gene expression were related to changes in mitochondrial DNA copy number, as we had observed in the muscle and PBMCs of patients previously (*46*), we calculated the mtDNA copy number in the bone and cartilage and observed no change in the human bone mtDNA copy number, but a decrease in the cartilage (**Figure 7G, H).** To validate the loss of the of the electronic transport chain (ETC) proteins, we performed a western blot of extracts from P14 bone and growth plate tissues. In the bone, we observed a decrease in complex I and Complex V (**Figure 7I-J)** and in the growth plate only in the Complex I (**Figure 7K).** We attempted to perform the same validation in the human bone samples but were not able to generate high quality protein extracts with the low amount of material we had on hand. Collectively, these data allow us to conclude that there is an alteration of mitochondrial processes, notably the loss of oxidative phosphorylation. In combination with our findings from the serum, we hypothesize that potentially the overburden and over import of cholesterol and fatty acids is causing a loss of the ETC complex members, as has been previously described (*89*).

**Figure 7:**
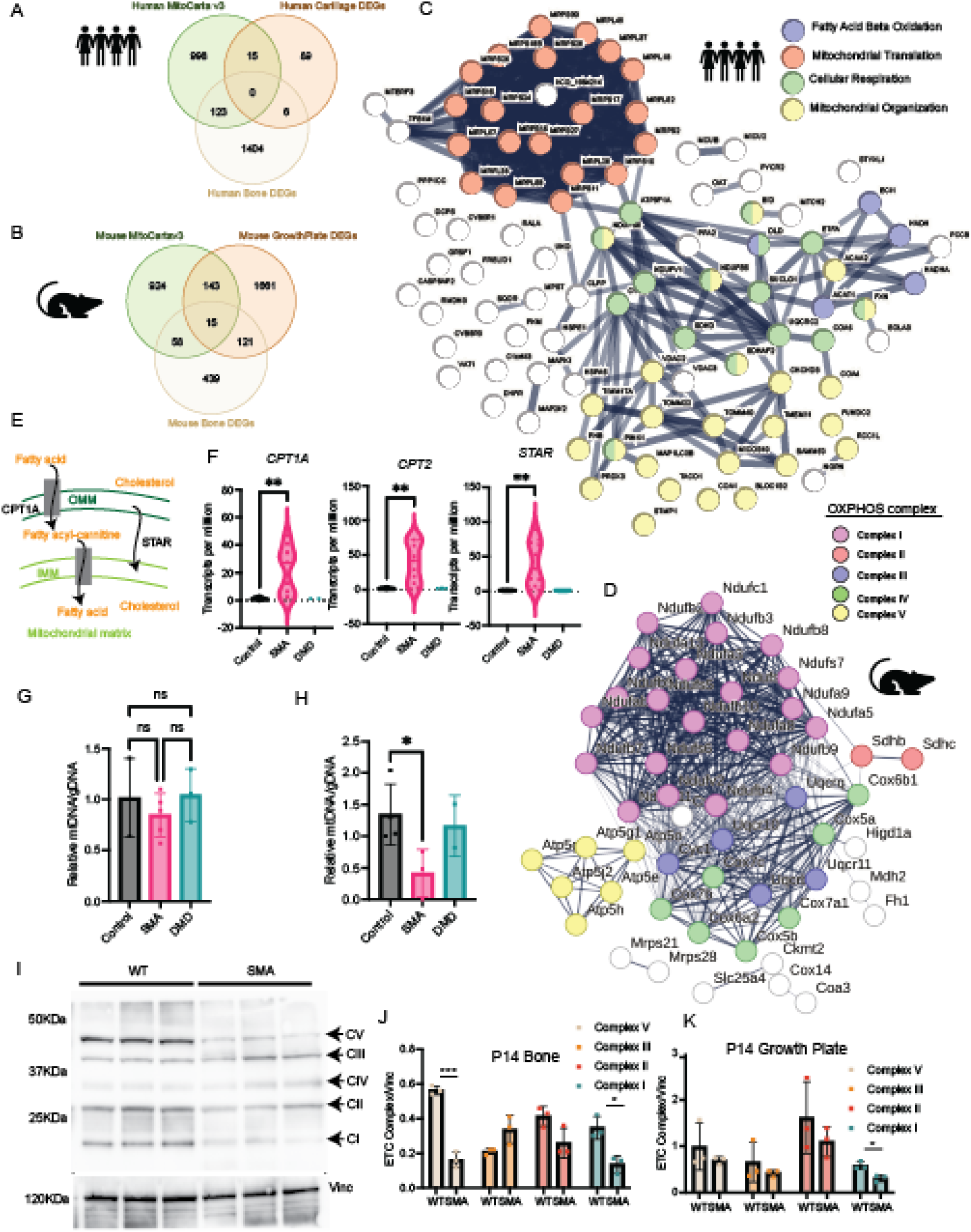
Mitochondrial OXPHOS changes are a common feature of Type II SMA patient and mouse model. **A.** Venn diagram overlapping the human MitoCarta 3.0 gene list with the differentially expressed genes in human SMA bone and cartilage. **B.** Venn diagram overlapping the mouse MitoCarta 3.0 gene list with the differentially expressed genes in mouse SMA bone and growth plate. **C.** STRING network analysis of the genes associated with the mitochondrial GO term in the downregulated DEGs in human SMA Type II vertebral bone. Each ball represents a single gene that is downregulated in SMA samples and is colored by its associated process. Nodes are connected by the known interactions, either physical or in the same network, that are known, and the thickness of the lines delineates the confidence of this interaction. **D.** STRING network analysis of the genes associated with the mitochondrial GO term in the downregulated DEGs in mouse at P14. Each ball represents a single gene that is downregulated in SMA samples and is colored by its associated process. Nodes are connected by the known interactions, either physical or in the same network, that are known, and the thickness of the lines delineates the confidence of this interaction. **E.** Schematic of the role of CPT1A, CPT2 and STAR in the import of fatty acids and cholesterol into the mitochondria. **F.** Violin plots of the transcripts per million (TPMs) of *CPT1A, CPT2* and *STAR.* Means are tested only between control and SMA samples using students t-test. ** < 0.001, DMD samples are plot as reference but were not statistically evaluated. **G.** Mitochondrial DNA (mtDNA) copy number relative to genomic DNA quantity in a subset of the SMA, DMD and AIS vertebral bone samples from the cohort. Means were tested by a one-way ANOVA followed by a multiple-comparison test. n.s – not significant. s that are found in the mitochondria and the DEGs from Type II SMA (from Grandi 2024) and in the Type II SMA vertebral bone. **H.** mtDNA copy number quantification in human cartilage **I**. Western blot of lysate of P14 *SMNΔ7* and WT mice bone tissue incubated with the anti-OXPHOS antibody cocktail and vinculin as the housekeeping gene. Each band represents an ETC complex, which is labeled. Complex IV was not detected in either blot. Complex I was not detected in the blot on the right. The proteins designating each complex are as follows: complex I, NDUFB8; complex II, SDHB; complex III, UQCRC2; complex IV, MTCO1; and complex V, ATP5A. **J.** Quantification of the level of each complex in P14 bone tissue from the western blot in J. **K.** Quantification of the level of each complex in P14 growth plate tissue.

### Irisin and other bone acting myokines are altered in SMA Type II patients

Muscle-bone crosstalk is an important part of the healthy maintenance of both tissues (*90*) and can be altered in a variety of disease states (**Figure 8A**). For example, recently it was shown that the gene expression changes in muscle associated with limb-girdle muscular dystrophy causes an alteration of the FNIP1-TFEB-IGF2 axis that affects bone metabolism (*91*). To determine if this alteration was occurring in our model systems, we began by performing RNA-sequencing of the *tibialis anterior* (TA) muscle of the P14 SMNΔ7 model to complement our RNA-sequencing of the hindlimb long-bones. We then assess the expression of 27 previously characterized myokines (*92*, *93*) in the SMNΔ7 and WT animals. Of these, 8 were downregulated in a statistically significant manner, including *Mstn, Sparc, Igf1, Fndc5, Ogn, Fstl1, Tgfb1,* and *Cxcl12* (**Figure 8B**). Changes in IGF1 have been previously documented in SMA mice and patients, with AAV mediated treatment with IGF1 prolonging lifespan in SMA mice(*94–97*). Given the importance of TGFB1 for cartilage and bone development, we measured serum levels in P14 *SMNΔ7* mice compared to controls, but did not observe any changes in the secreted levels, suggesting that any alterations in production in the muscle were localized (**Supplemental Figure 5A**).

**Figure 8:**
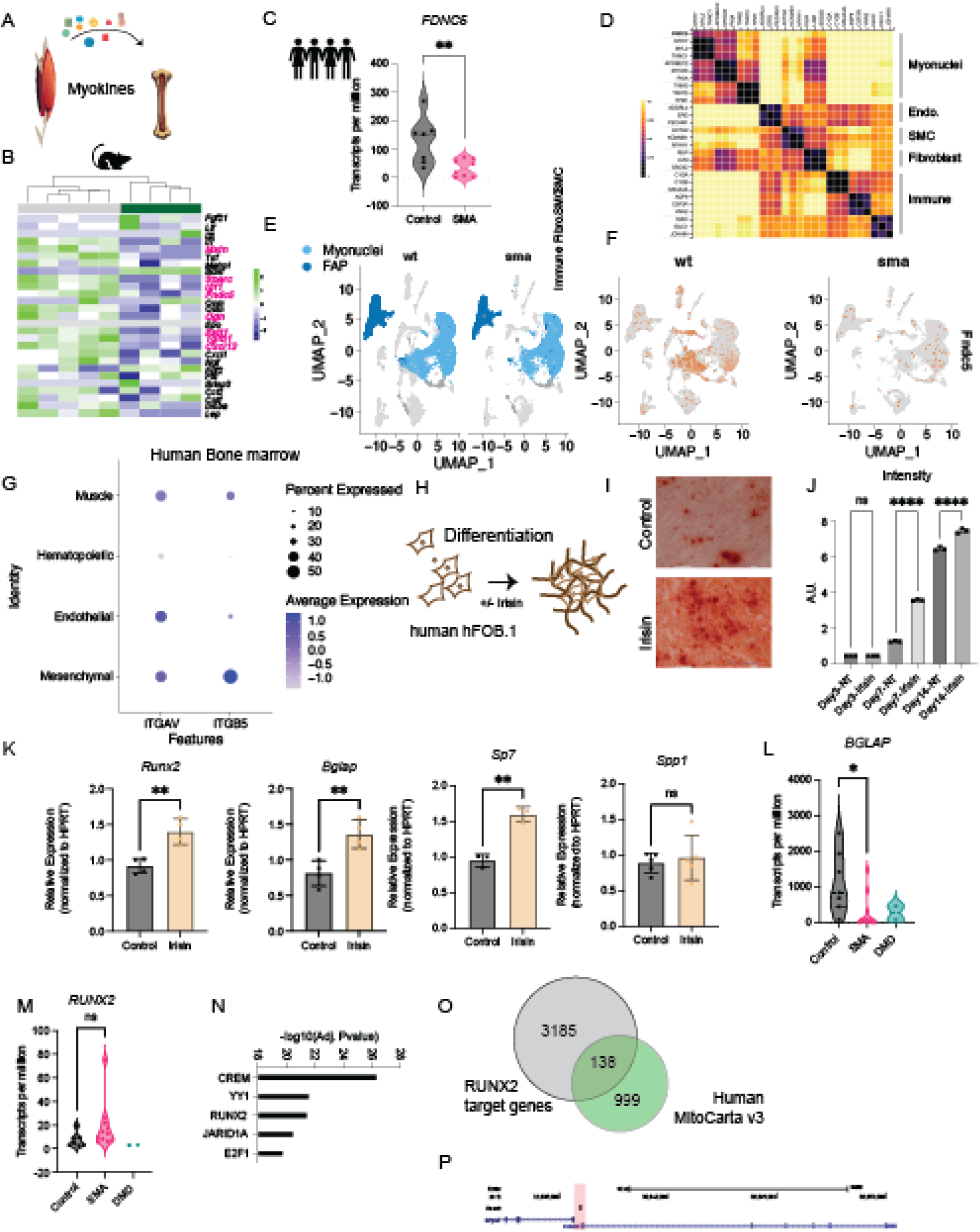
Muscle-to-bone signaling via Irisin is altered in SMA. **A.** Schematic of myokines communication to bone. **B.** Heatmap of selected myokines with known roles in bone biology. Those highlighted in pink are differentially expressed in the TA of *SMNΔ7* compared to WT controls. **C.** Violin plot of the transcripts per million (TPMs) of *FDNC5* (Irisin). Means are tested only between control and SMA samples using students t-test. ** < 0.001, DMD samples are plot as reference but were not statistically evaluated. **D.** snRNA-seq reanalyzed from the Human Protein Atlas for the expression of Irisin (*FNDC5)* versus other cellular markers. **E.** UMAPs of single-nucleus (sn) RNA-sequencing of the tibialis anterior of *SMNΔ7* mice at P14 compared to WT controls. Myonuclei are highlighted in light blue and fibroadipogenic progenitors (FAPs) in dark blue. Each point represents a single nucleus that was captured **F.** Same UMAP representation as in E, but now each point is colored by the expression of *Fndc5*. **G.** Expression of the Irisin receptors *ITGAV* and *ITGB5* in human bone-marrow data derived from study QQQ. **H.** Schematic of the differentiation experiment with irisin treatment in the human immortalized osteoblast cell line hFOB.1. **I.** Representative alizarin red staining images from osteoblast treated with irisin for 7 days or with vehicle control. **J.** Quantification of alizarin red staining at Day 3, 7 and 14 of differentiation. K. qRT-PCR of the expression of osteoblast genes in hFOB.1 cells treated with irisin or vehicle control for 72 hours. **L.** Violin plot of the transcripts per million (TPMs) of *BGLAP* in human bone samples. Means are tested only between control and SMA samples using students t-test. ** < 0.001, DMD samples are plot as reference but were not statistically evaluated. BM. Violin plot of the transcripts per million (TPMs) of *RUNX2* in human bone samples. Means are tested only between control and SMA samples using students t-test. ** < 0.001, DMD samples are plot as reference but were not statistically evaluated. **N.** Top transcription factor motifs that were over-represented in the downregulated genes in SMA bone samples. **O.** Overlap between the RUNX2 target genes derived from EnrichR and the human MitoCarta (*87*) genes. **P.** Representative capture of Runx2 Chip-seq studies in mouse osteoblasts(*117*) showing RUNX2 binding at the promoter of the genes *Ndufb5*.

To test if the same changes in myokines were present in the SMA Type II patients, we mined our previously published cohort of 8 sample Type II paravertebral muscle samples (*46*). Unlike in the mouse samples, there was not a clear clustering of samples by the expression of the myokines, suggesting that compared to the AIS controls, there is not a global alteration in myokine production (**Supplemental Figure 5B**). However, among all the myokines, there was a statistically significant increase in the amount of *IL6* (**Supplemental Figure 5C**) and a decrease in *FNDC5*, also known as Irisin, mimic the results in the mouse (**Figure 8C**). To better understand which cells in the muscle expressed Irisin, we first mined previously published snRNA-sequencing maps of human muscle (*98*, *99*) and found *FNDC5* expression is highest in myonuclei (**Figure 8D**). To validate these findings in our mouse model of SMA, we generate single-nucleus RNA-sequencing data (snRNA-seq) from the TA of a P14 SMA mouse and compared it to a previously existing ENCODE P14 dataset (*100*) (**Figure 8E**, **Supplemental Figure 5 D-F**). As expected, we found a loss in the population of myonuclei found in SMNΔ7 mice, including the loss of myonuclei expressing *Fndc5* and other key myokines (**Figure 4F, Supplemental Figure 5D, G**).

We next wanted to understand which cells in the bone and bone marrow niche can receive the irisin signals. Using the published map of the human bone marrow niche, we looked for the two known receptors of Irisin, ITGAV and ITGB5 (*101*, *102*). The single cell data showed that while ITGAV expression is ubiquitous among the BM populations, the ITGB5 is more restricted to mesenchymal populations (**Figure 8G**). To assess the role that this could have on human osteoblast differentiation, we treated a human osteoblast cell line hFOB.1 with *Irisin* during differentiation (**Figure 8H**). Given the anabolic role of Irisin on the bone (*103*), we analyzed the calcium deposition in cells treated with irisin and controls after 3, 7 or 14 days of culture. Alizarin red staining was increased at 7 and 14 days (**Figure 8I-J, Supplemental Figure 5H**). We next assessed the gene expression profiles at 72 hours post treatment (day 3), reasoning that irisin should provide a post in pro-osteogenic factors. Indeed, we observed the upregulation of *Runx2*, *Bglap (osteocalcin) and Sp7,* but not of *Spp1* (**Figure 8K**). Among these targets, *Bglap* is downregulated in the human vertebral bone transcriptomes of SMA patients (**Figure 8L**). Although *Runx2* itself is not downregulated (**Figure 8M),** several of its target genes are downregulated, with the RUNX2 motif being the third most enrichened motif among the downregulated genes in SMA bone samples (**Figure 8N**). RUNX2 is known to be responsible for many pro-osteogenic gene process and additionally has known binding sites in many of the genes controlling the mitochondrial processes (**Figure 8O**) that are altered in the SMA bone. This includes, for example, *NdufB5*, a common target between the mitochondrial genes in human and mouse samples (**Figure 8P**). As such, irisin may provide an interesting target for rebalancing the pro-osteogenic environment in the bone (**Figure 5**) and boosting expression of mitochondrial genes (**Figure 7**).

## DISCUSSION

In recent years, the multi-organ nature of spinal muscular atrophy (SMA) has become evident (*3*). Among the affected tissues is the skeletal system, as demonstrated both by mouse models (*32*, *34*, *35*) and by the fracture and scoliosis symptoms in treated SMA patients (*29–31*). Here, we present, to our knowledge, the first transcriptomic characterization of human Type II SMA vertebral bone and cartilage tissues and compare the observed changes to our mouse model of SMA, the SMNΔ7 model. Although the mouse and human samples have many differences, we found common changes in the regulation of endosomal membrane pathways, fatty oxidation, mitochondrial OXPHOS and changes in the signaling of PPARγ and several myokines including Irisin. Our data highlights that, in addition to the cell-intrinsic features that have been shown by other studies, alterations in SMA bone health are the consequence of multi-systematic factors, originating from various organ systems, including the liver and the muscle. By characterizing these molecular changes, we set the stage for the implementation of targeted combination therapies that can address these underlying bone health issues, in addition to the current standard of care. Additionally, this study highlights in general the multisystemic nature of bone health, regardless of an underlying NMD.

### Effects of current treatments on SMA bone health

Using our previously published AAV9-PGK1-SmnOpti gene therapy (*57*, *58*), we were able to test the hypothetical scenario of a highly effective gene therapy given at post-natal day 1. Although this mirrors some aspects of treatment with Zolgesma, it is the best-case scenario, as pup are treated before the onset of symptoms and with an administration route that is significantly more efficient than the intravenous injection used in the clinics (*58*). We showed that this dose of gene therapy successfully transduces all the organ systems, with variable VGCNs, and can deliver some SMN protein to all tissues, although in many it does not achieve the wildtype levels. Notably, these P1 injections result in the transduction of cartilage and bone at PND7, although with viral copies per genome lower than in many other tissues, and at PND7 the animals express SMN protein. These VGCN copies are lost with time, with very few SMN RNA from the AAV present at P14 and essentially no RNA or viral copies at 6-months. This corresponds to the total absence of SMN protein observed in the 6-month inject samples. Despite, this, the bone can continue to function normally, with no alterations overserved both at the whole-skeletal mount level and more important, a full restoration of the transcriptomes at P14 and 6-months post injection. This suggests that regardless of the need for SMN in the cells early on, the bone tissue can survive postnatal life without SMN protein. We suggest that this brings further evidence to the hypothesis that bone health in SMA patients is primarily driven by the systemic changes, including changes in liver, fatty acid and cholesterol metabolism and muscle health, which affect the bone. In patient samples, while we could not measure the protein levels of SMN due to technical problems, we did not observe the same amount of transcription of full-length SMN from the *SMN2* copies to compensate for the loss of SMN1, in contrast to what we observed in the muscle. However, our findings from the mouse suggest that delivering a copy of SMN is not of primary importance in the long-term, and that combination therapy efforts should be focused on managing the muscle symptoms and the metabolic disease.

These findings further support the notion that simply restoring SMN or improving muscle function may not be sufficient to fully rescue bone integrity or prevent skeletal deformities in SMA. While existing treatments, such as bisphosphonates, primarily address bone density loss secondary to immobility, they do not target the underlying molecular disruptions intrinsic to SMA-affected bone. Therefore, a deeper understanding of the molecular mechanisms driving bone pathology in SMA is critical for developing novel or adjunctive therapies that directly improve skeletal outcomes to enhance overall patient quality of life.

### Effects of increased cholesterol and fatty acids on bone health

Cholesterol and fatty acid metabolism have complex relationship with bone health, with both positive and negative outcomes being attributed, depending on the type of lipid, the dosing and the timing (*73–75*). As for many other tissues, lipids present a key source of nutrients for the bone. Additionally, cholesterol is important for maintaining membrane compositions, particularly in osteoclasts, which have been shown to require particularly high levels of cholesterol to maintain their unique membrane properties that allow for digestion of the bone matrix (*77*, *78*). However, it has been shown in the muscle and in the liver, an overabundance of either fatty acids or cholesterol can overwhelm mitochondrial processes, leading to a loss of oxidative phosphorylation and ATP production(*89*, *104*). Collectively, our data suggests a model where due to hereby uncharacterized roles of SMN in the liver, there is an accumulation of cholesterol and fatty acids both in the liver, and other tissues such as the skeletal muscle and the spinal cord. Indeed, this overload of cholesterol and fatty acids can be transmitted to osteoblasts via the serum of symptomatic animals and activates PPARG signaling, the same signaling pathway that is present both in our mouse and human data. It is possible that this overload also leads to the loss of mitochondrial function seen both in the mouse and human data. Importantly, this defect can be corrected by our gene therapy, which decreases the cholesterol accumulation in the liver after delivery of high levels of SMN. This data suggests that drugs which target cholesterol and fatty acid metabolism, such as statins, could be an interesting target for combination therapies in SMA patients, and that this may have whole-body effects, including in the bone. Targeting cholesterol metabolism has also been suggested a potential therapeutic avenue in Duchenne muscular dystrophy, from findings in the skeletal muscle(*105*).

### The role of muscle to bone signaling in SMA bone health

Bone and muscle have a tightly connected relationship, with both mechanical and biochemical cues being exchanged between the two tissues. As such, it is likely that the increased incidence of scoliosis observed in the SMA Type II patients is due in part to the weakness of the paravertebral muscles, which in turns stems from their denervation (*46*). Muscle changes have been observed in idiopathic scoliosis cases, independent of NMD (*106–109*). Due to this tightly interlaced relationship, we used our previously published dataset on Type II paravertebral muscle to investigate changes in secreted myokines with known osteogenic effects. Among these, in the human data we observed changes in IL6 and Irisin. Both myokines have been associated with exercise, with Irisin being increased by physical activity and IL6 being lowered. As such, we could hypothesize that their modulation in the opposite direction as what is expected after exercise, is due to the limited mobility of SMA Type II patients. Regardless of the mechanisms, the alteration of these myokines can have profound effects on bone health. For example, IL6 has been found to have roles in the indirect stimulation of osteoclast formation, and the IL6 family of receptors is expressed in osteoblast cell lineages (*110*, *111*). Although we did not test the effect of IL6 on osteoclast formation directly, its pro-osteoclastogenic effects would be in-line with our findings that SMA Type II patients have an enriched osteoclastogenic signature. Irisin also has both pro-osteogenic and pro-osteoclastogenic properties but is generally accepted as helping to maintain the balance between the two processes. Our data demonstrates that Irisin can effectively increase pro-osteoclastogenic gene expression, and previous Chip-seq experiments have shown that it can target a variety of mitochondrial genes as well, making it an attractive putative combination therapy, as we observed many changes in mitochondrial pathways.

### Limitations of the study

Our study suffers from the problems common to studies attempting to characterize changes in tissues of living patients with a rare disease – material is difficult to obtain, highly heterogenous and not present in large quantities. Despite these limitations, we were able to obtain a cohort of 11 SMA vertebral bone samples, where 8 of the 11 samples SMA had similar expression patterns. Unfortunately, due to our limited ability to have full clinical histories for these samples, we cannot exclude effects due to disease onset or time of treatment that may explain these differences. Additionally, our study was not powered enough to assess the effect of sex, if any, on the severity of the bone changes observed in SMA. Larger cohort studies will be necessary to fully delineate the natural history of bone in SMA patients. However, this study provides the first molecular map of the changes occurring in patients, and suggests putative pathways, including improving fatty acid and cholesterol metabolism and increasing the myokine Irisin, which could be used for putative combination therapies.

## MATERIALS & METHODS

### Sex as a biological variable

In this study, biological sex taken into consideration when we compared the male and female sample in our cohort. We explicitly tested for differences in the SMA presentation based on sex, but no correlation was found between SMA subgroups for sex, and no sex effect was found among the differentially expressed genes. However, due to limited sample availability, we did not have a balanced cohort (n=2 females, n=9 male SMA Type II samples) thus this study is now power enough to discover sex differences in SMA Type II vertebral bone. For mouse samples, sex was not considered as we have not observed any difference in survival between the male and female mice.

### Study approval

Samples were obtained with patient or parental written consent, collected from the surgical residue from patients undergoing surgery for spinal sclerosis. Samples were collected under the authorization of the Minister of Research, approval number AC-2019-3502, in accordance with French and European laws.

### DNA extraction from human tissues

DNA was extracted from approximately 2–3 mg of frozen muscle tissue using the Qiagen DNeasy Blood and Tissue Kit. Samples were first ground into a powered using a tissue grinder and then digested for 3–6 hours with proteinase K, as per the manufacturer’s recommendations.

### RNA extraction from tissues

Frozen samples (∼3–4 mg) were ground in a tissue grinder, making sure to keep the tissue frozen. The samples were then incubated with 500 μL of Trizol and ceramic beads and the tissue was homogenized using a tissue lyzer until no tissue was visible. Samples were spun down to remove any remaining undigested tissue. RNA was then extracted by adding chloroform and taking the aqueous phase. Extracted samples were cleaned using the Qiagen RNeasy Mini Kit. RNA quality was assessed using an Agilent TapeStation. All samples had an RIN of greater than 5. For cells, 500,000 cells were collected by spinning and resuspended in RLT buffer according to the manufacturer’s specifications and directly extracted using the Qiagen RNeasy Mini Kit.

### cDNA library generation and sequencing

Libraries for sequencing were prepared using the SMART-Seq v4 PLUS Kit (Takara/CloneTech) according to the manufacturer’s specifications, using 10 ng of input RNA. Samples were sequenced after barcoding using the Illumina NovaX platform with 150-bp paired-end reads.

### Library alignment and differential gene expression analysis

Fastq files were analyzed using the nf-core RNA-seq pipeline with the standard parameters (83). Reads were aligned to hg38. Differential expression analysis was performed using DESeq2 (84), and R was used for visualization of the data. Pathway analysis was performed using EnrichR (85) and Metascape (86).

### Mouse husbandry

Animals were bred and housed as previously described (*58*) under controlled conditions following the French and European guidelines for the use of animal models (2010/63/EU). Briefly, breeding pairs of triple transgenic (*SMN2^+/+^, SMNΔ7*^+/+^, *Smn*^+/−^; no. SN 5025) were purchased from The Jackson Laboratory. The genotype of the offspring can be WT (*SMN2*^+/+^, *SMNΔ7*^+/+^, *Smn*^+/+^), heterozygous (*SMN2*^+/+^, *SMNΔ7*^+/+^, *Smn*^+/−^), or knockout, named SMNΔ7 (*SMN2*^+/+^, *SMNΔ7*^+/+^, *Smn*^−/−^). Mice were genotyped using the previously established primers. Mice were euthanized 14 days post-injection for SMN expression analyses by intraperitoneal anesthetic injection (10 mg/kg xylazine, 100 mg/kg ketamine), followed by intracardiac perfusion, with PBS. Euthanasia was performed according to regulations and tissue samples were recovered and immediately frozen in liquid nitrogen for further downstream processing, depending on the experiment. For the embryonic samples, pregnant females were euthanized at E17.5. All animals were approved by the French Ministry of Higher Education and Research and humane endpoints established in the animal protocol (Agreement number #19475 and #46937).

### AAV Vector Production and Titration

The AAV expression cassette was produced as previously described (*57*, *58*). Briefly, the expression sequence for codon-optimized human *SMN1* of the ubiquitous PGK promoter was cloned by enzymatic restriction into the previously described deleted ITR2 of the AAV genome. The serotype 9 AAV vectors were produced using helper virus-free transient transfection in HEK293 cells following the protocol described by the MyoVector Platform. Cells are transfected using two helper plasmids and the pAAV transgene using suspension culture of HEK293T cells. After cell lysis, suspensions are centrifuged and purified on an iodixanol gradient. Finally, the AAV vector is concentrated and resuspended in DPBS. in suspensions HEK293T cells Each production was quantified by real-time PCR and vector titers were expressed as vg/mL.

### *In Vivo* AAV Injection

Injections of new-born mice (post-natal day 1) were performed using a 50-μL Hamilton syringe with a 31G and 30-mm length needle. 7 μL of viral suspension, containing 5 × 10^10^ vg was administered into the brain lateral ventricle through a unilateral i.c.v. injection (1 mm anterior, ±1 mm lateral to the lambda, and 2 mm deep).

### DNA extraction from mouse tissues for viral copy number measurement

For tissues from P3 and P7 and P14 animals, DNA was extracted as for the human tissue samples described above. For tissues derived from long-term animals, we used the Qiagen Puregene core kit A according to the manufacturer’s instructions due to difficulties precipitating the protein components using the other kit.

### Viral copies per genome (VGCN) measurement in injected mouse tissues

DNA samples were extracted as above and diluted to a concentration of 10ng/ul in water. VGCN was measured using a multiplex Taqman quantitative qPCR system using *Ttn*-VIC probes to determine the copies of nuclear DNA and *SV40*-FAM to determine the number of AAV copies. Reactions were carried out using the Applied Biosystems Taqman master mix in a volume of 10 ul with 4 ul of DNA at 10 ng/ul per reaction on an Applied Biosystem’s QuantStudio™ 7 Pro with the following cycling conditions: 95° C for 10 min and 40 cycles of 95C for 15 sec, 60°C for 1 min. Viral genome copy number values per were calculated using a standard curve derived from a plasmid that contains both primers to obtain the number of copies of each genes and dividing the viral copy number by the housekeeping gene copy number to express VGCN by genome. The probe for SV40 is AGCATTTTTTTCACTGCATTCTAGTTGTGGTTTGTC with the forward primer: AGCAATAGCATCACAAATTTCACAA and the reverse primer: CCAGACATGATAAGATACATTGATGAGTT and for *Titine*, the probe sequence is: TGCACGGAAGCGTCTCGTCTCAGTC the forward primer: AAAACGAGCAGTGACGTGAGC and the reverse primer: TTCAGTCATGCTGCTAGCGC.

### Whole Mount Skeletal Staining

Post natal day 3, 7 and 14 pups were euthanized according to the established protocols and the skin and muscle, and internal organs were removed. Whole mount skeletal staining was performed as previously established(*112*). Samples were destained in 1% KO/20% glycerol for several weeks to remove the extra dye. Images were taken with a dissection microscope.

### Protein extraction and Western blotting

Protein extracts were prepared from liquid nitrogen–frozen human muscle tissues. Tissues were lysed in RIPA buffer (Sigma-Aldrich) supplemented with a protease inhibitor cocktail (Complete Mini, Roche Diagnostics). Proteins were quantified using the DC Protein Assay (Bio-Rad) and 40 μg of protein was loaded. In all blots except those using the mitochondrial complex cocktail enzyme, proteins were heated to 100°C for 5 minutes; proteins used for mitochondrial OXPHOS cocktail blots were heated to 37°C. Proteins were run in 4%–10% Bis-Tris gels (Bio-Rad) and transferred to membranes using the Turbo Transfer System (Bio-Rad). The membranes were then incubated with the corresponding antibody: anti-SMN (1:1,000; BD Biosciences, 610647), anti-vinculin (1:1,000; Sigma-Aldrich, V9191), or anti-mito cocktail (1:500; Abcam, ab110413) diluted in Tris-buffered saline containing 0.2% Tween 20 (TBST, Bio-Rad) supplemented with 5% nonfat dry milk. Primary antibody incubation was performed overnight. After 3 washes in TBST, the membranes were incubated with a horseradish peroxidase–conjugated secondary antibody (1:10,000; GE Healthcare) diluted in the blocking buffer. The membranes were further developed using the chemiluminescence substrate SuperSignal Ultra reagent (Thermo Fisher Scientific) or the SuperSignal West Femto Maximum Sensitivity Substrate (Thermo Fisher Scientific) in the case of SMN and imaged on a Bio-Rad ChemiDoc MP imaging system.

### Alizarin red staining of osteoblasts and quantification

Osteoblasts were grown in 60 mm plates. At the desired time point, cells were washed three times with PBS and fixed with 4% PFA at room temperature for 15 minutes, followed by three washes in PBS. Fixed plates were stored with PBS at 4C until all time points were collected. Before staining, plates were washed one time with distilled water for 1 minute. 40mM Alizarin Red solution (REF) was added to the plates for 30 minutes at room temperature with gentle rocking. After 40 minutes, cells were washed three times in PBS before imaging on the Keyance microscope.

For quantification, 1mL of 10x acetic acid solution in PBS was added to each plate and plates were incubated at RT for 30 minutes to lift the coloration. The solution was then collected in a tube and heated to 85C for 10 minutes, placed on ice and once cooled, centrifuged for 5 minutes at 20,000 X G. The supernatant was transferred to a new tube, taking care not to bring any precipitates that formed after spinning. The pH was balanced using 50mM NaOH (200 ul per 800 ul of acetic acid solution). Absorbance was read on a 96-well flatbottom plate at 405nM.

### Real Time quantitative -PCR

cDNA was generated using the Applied Biosystems High Capacity Retrotranscriptase Kit (ThermoFisher) using between 1-2 ug of RNA, depending on the experiment. Each experiment also contained one RT-control to measure the amount of contaminating gDNA, if any. After retrotranscription, samples were diluted to a total of 40 ul and 2 ul was used to carry out reactions with either SybrGreen (Applied Biosystems) or Taqman (AppliedBiosystems) reactions. A list of primers used can be found in Supplemental Table 1.

### Isolation of Nuclei from Mouse Muscle

Nuclei were isolated from the gastrocnemius muscle of a postnatal day 14 SMA mouse. The muscle was cut using a cryostat into 10 slices of 50uM. These slices were then used for nuclei isolation using dounce homogenizers. Same homogenization buffer was prepared as previously described(*113*), with minor modifications. Briefly, the homogenization buffer consists of 250mM sucrose, 25mM KCl, 5mM MgCl2, 20mM Tricine-KOH (ph 7.8) with 1mM DTT, 0.5mM spermidine, 0.15mM spermine, 0.3% NP40 and complete Protase inhitor (Roche) added on the day of extraction. Homogenization buffer was spiked with 2% BSA to prevent nuclei from sticking to each other and 5ul of RNAseinhibitor (Promega, 40U/uL) were added to 1 mL of buffer in the dounce homogenizer prior to isolation. Muscles cuts were dounced 10 times with pestle A and 8 times with pestle B. The lysate was then filtered through a 100uM bucket filter, before being filtered through a 70uM and then 30uM flow-mi filter. Nuclei were spun down for 10 min at 500 rcf at 4C and resuspended in nuclei buffer (10x Genomics) with DTT and RNAse inhibitor and counted using DAPI. 25,000 nuclei were loaded into the 10x Genomics 3’ scRNA-sequencing reaction as per the manufacture’s instructions.

### 10x Genomics single-nuclei RNA-sequencing library preparation

10x single-nuclei libraries were prepared with the 3’ Gene Expression kits from 10x using the v3.1 chemistry and dual-indexes following the manufacture’s protocol. The sample, pooled with other libraries not included in this work, was sequenced using an Illumina NovaX instrument according to the manufacture’s instructions.

### Analysis of single-nucleus RNA-sequencing libraries

Data was aligned to the mm10 mouse genome reference from 10x genomics using CellRanger (version 7.1.0). The raw and filtered outputs were then used for downstream analysis in R. First, SoupX (*114*) was used to remove ambient RNA from the samples. Doublets were then called using DoubletFinder(*115*). These modified matrixes were then used to generate Seurat objects for clustering and analysis. The SMA P14 gastrocnemius was merged with existing data on wildtype P14 gastrocnemius snRNA-seq from ENCODE (Accession ENCD0509HIY; https://www.encodeproject.org/biosamples/ENCBS759QMD/). Data was merged using Seurat’s CCA method.

### Reuse of single-nucleus RNA-sequencing libraries

A publicly available single-cell RNA-seq dataset of the human bone marrow was utilized to perform deconvolution of the bulk vertebral bone transcriptomes. Note that bone marrow atlas was not derived from vertebral bone but from the femoral head of patients undergoing total hip arthroplasty. The Seurat object including its annotations were downloaded from GEO (GSE253355) and the authors original annotations were used.

### Deconvolution of bulk RNA-sequencing transcriptomes

Bulk RNA-sequencing transcriptomes from the vertebral bone were deconvoluted using the granulator package(*44*), using the dtangle method (*45*). As granulator easily allows you to compare between multiple methods, other methods were compared but not significant changes in estimation were observed.

### Staining mouse tissues with Filipin III and Sudan Black

10µm-thick liver cryosections were fixed 10 minutes with paraformaldehyde, quenched with 1,5mg/ml glycine in PBS for 10 minutes, then stained with Filipin III (SAE0087, Sigma Aldrich) at 250µg/ml for 2h, in the dark, before washing in PBS. Slides were mounted with PBS.

For Sudan Black staining, liver cryosections were washed with deionized water, dehydrated with 100% propylene glycol for 5 minutes, and then stained with Sudan Black B (199664, Sigma Aldrich) for 2h before washing in deionized water and differentiation in 85% propylene glycol for 3 minutes. Slides were then mounted in aqueous mounting medium.

Staining was imaged with a Keyence VHX-7000N. All histological analyses were performed in a blinded manner on Image J.

### Measurement of cholesterol and other metabolic products of cholesterol by mass-spectrometry

Samples were weighted and subjected to a double lipidic extraction followed by an alkaline hydrolysis of the samples 1h at 60°C. Sterols and oxysterols were separated by thin-layer chromatography and derivatization of the samples were done with BSTFA/TMCS 1% before injection in GC-MS/MS (Agilent 7890B/7000C). Cholesterol 3C13 and 7β-hydroxycholesterol-d7 were used as internal standards.

### In vitro culture and differentiation of human osteoblast cell line

hFOB1.19 (CRL-3602) cells were obtained from ATCC. Cells were cultured in a 1:1 mixture of DMEM: F12 (QQQ), and DMEM High Glucose (QQQ) with 1x Glutamine and 10% FBS (Corning) and passed with 0.05% trypsin when they reached 70% confluency. Cells were differentiated DMEM augment with 10% FBS, 5mMBeta-glycerol phosphate, 50mg/mL ascorbic acid, 10uM Dexamethanone, and 4mM L-glutamine.

### In vitro treatment of osteoblasts with Irisin

Osteoblasts in differentiation medium were treated with 10 ng/mL Irisin (Peprotech) for either 72hours for RNA experiments or 3, 7 and 14 several days for staining experiments. Media was changed every 24 hours during treatment.

### In vitro treatment of osteoblasts with mouse serum and staining with Fillipin III

Osteoblasts in differentiation medium were treated with differentiation media with 10% mouse serum and 1x gentamicin (Corning). For all samples, the serum for 3-4 mice was pooled and mixed. Samples were cultured for 72 hours, and media with new serum, from the same pooling batch, was changed every 24 hours.

### In vitro differentiation of osteoclasts

Osteoclasts were isolated from mice according to a previously established protocol, with some modifications(*116*). Postnatal day 14 mice (WT or SMND7) were sacrificed according to previously established protocols and both front and hindlimb bones were dissected and cleaned. The ends were cut, and cells were flushed two times with cold basal RPMI media (RPMI + 10% FBS) and antibiotics. Cells were then spun down and resuspended in 1x red-cell lysis buffer (QQQ) and incubated at room temp for 8 minutes. An equal volume of RPMI was added to quench the reaction and cells were spun down. Cells were then placed in culture at 100,000 per well of a 24-well plate and treated for 3 days with 30ng/mL of MCSF (QQQ). On Day 3, cells were changed to media containing 30 ng/mL of MCSF and 100ng/mL of RANKL. Media was changed every 2 days. Cells were collected for TRAP staining or RNA isolation on Day 8.

### TRAP staining

Osteoclasts were differentiated on IBIS chambered slide plates and stained using the Leukocyte TRAP staining kit (Sigma) according to the manufacturer’s instructions.

### Data availability

Raw RNA-seq fastq files are available in the NCBI GEO under accession number GSE280223.

## Acknowledgements

We thank the patients and their families for consenting to the donation of tissue samples to this research. We would also like to thank the members of the Smeriglio lab for their helpful discussions and the MyoBank platform which allows for the collection of human samples, as well as MyoImage, which supports the microscopy center and maintains the Keyance microscope, and the MyoVector Platform that supports the production of the AAV. We would like to thank the staff of IGenSeq Platform (Institut du Cerveau) for their help with the RNA-seq library quality control and use of their facilities. We would like to acknowledge the Plateforme de lipidomique fonctionnelle (IBiSA: Infrastructure en Biologie Santé et Agronomie) associated with l’IMBL (Institut Multidisciplinaire de Biochimie des Lipides) for the lipidomics measurements in the muscle and spinal cord. We would like to acknowledge the Institut Francais de Bioinformatique (IFB), which provides the HPC server used for the RNA-seq analysis. We thank the Penn Vector Core, Gene Therapy Program, University of Pennsylvania, Philadelphia for providing the pAAV9 plasmid (p5E18-VD29). This work was supported by the Association Française contre les Myopathies (AFM), the Association Institut de Myologie (AIM), the Sorbonne Université, and Fondation Carrefour. This work was also supported by the Fondation Maladies Rares GenOmics (2023) (grant 041703). FCG was financed through support of the France Relance National Program and the Marie Curie Postdoctoral Fellowship (GAP: 101109098) and Institut National de la Santé et de la Recherche Médicale (INSERM), PS is funded through support of the Institut National de la Santé et de la Recherche Médicale (INSERM), AA was financed through the support of the Region Ile-de-France in the framework of DIM doctoral fellowship, EG was financed through support of the Eramus+ Scholarship 2023 and the Sorbonne Universite ED515 doctoral fellowship, YS was financed through French Committee for the Evaluation of University Cooperation with Brazil Program (CAPES Cofecub) N° 32/2022.

---

**Supplemental Figure 1:**
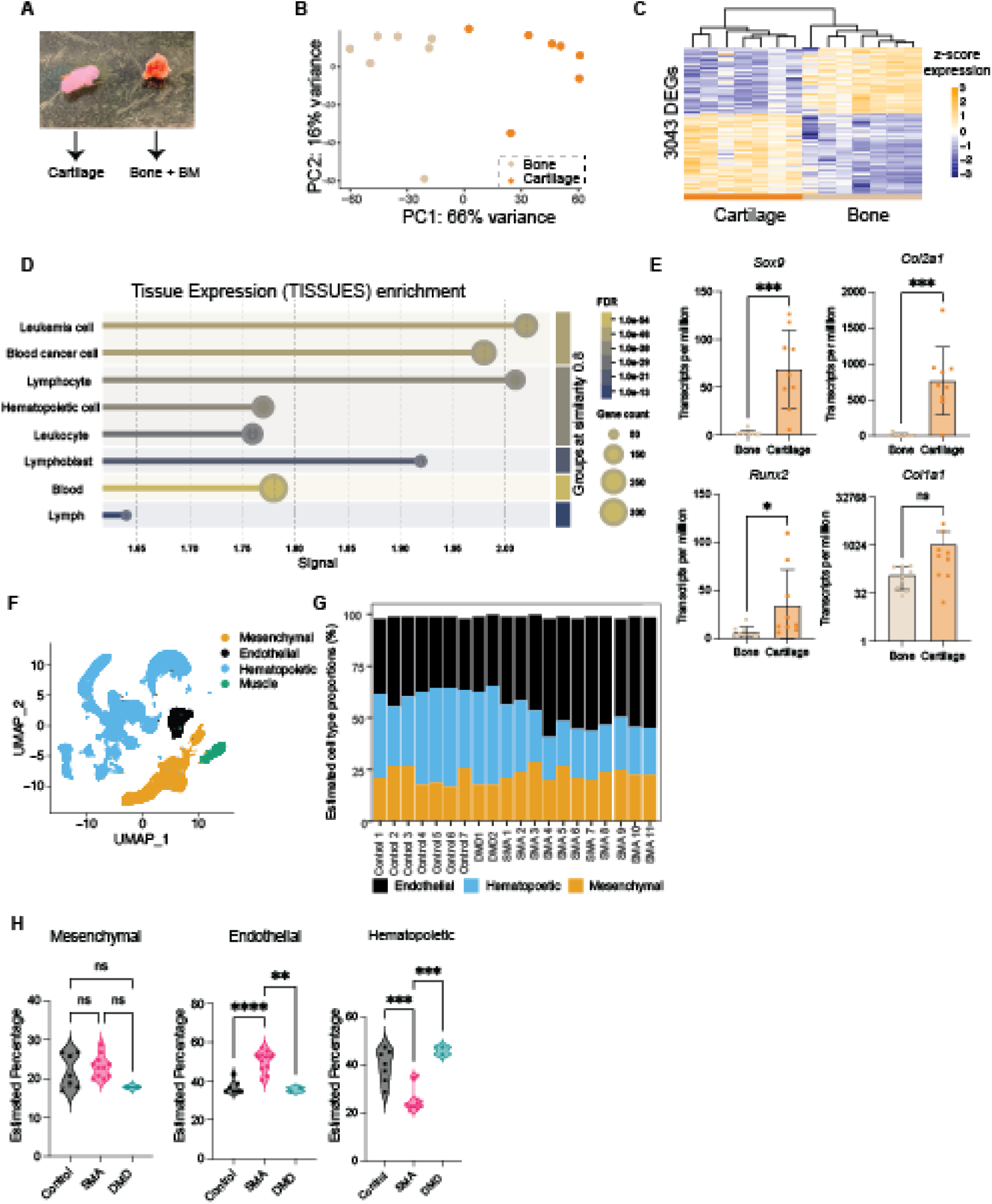
Characterization of the transcriptome of human vertebral bone and cartilage from SMA, DMD and AIS patients. **A.** Representative image of the cartilage and bone pieces recovered from scoliosis surgery. Samples were dissected to remove any additional connective tissue or fat, washed several times in sterile PBS and then snap frozen for molecular biology. **B.** Principal component analysis (PCA) plot of the transcriptome of the AIS control cartilage and bone samples compared to each other. Each point represents a single sample. Tissue type represents 66% of the observed variance in the transcriptomes. **C.** Heatmap of the 3034 differentially expressed genes between AIS control cartilage and bone. Unsupervised hierarchical clustering is shown at the top. The gene expression is z-scored across each row. **D.** Tissue expression enrichment calculated using EnrichR on the genes upregulated in the bone samples. **E.** Plots of the transcripts per million (TPM) of the chondrogenic genes *Sox9* and *Col2a1* and the osteogenic genes *Runx2* and *Col1a1*. The average expression between each tissue was compared using a student’s t-test. * < 0.01, *** <0.001. **F.** Single-cell RNA-sequencing map of human bone marrow taken from dataset GSE253355, with the authors original annotations (*43*). **G.** Estimated percentage of each cell type for each bone transcriptome based on the single-cell RNA-seq data in (F). Note that the muscle subtype was not predicted. **H.** Estimated percentages as plotted in panel F represented as violin plots divide by patient group. The average between each group was tested using a one-way ANOVA with multiple comparisons. * < 0.01, *** <0.001, **** <0.0001

**Supplemental Figure 2:**
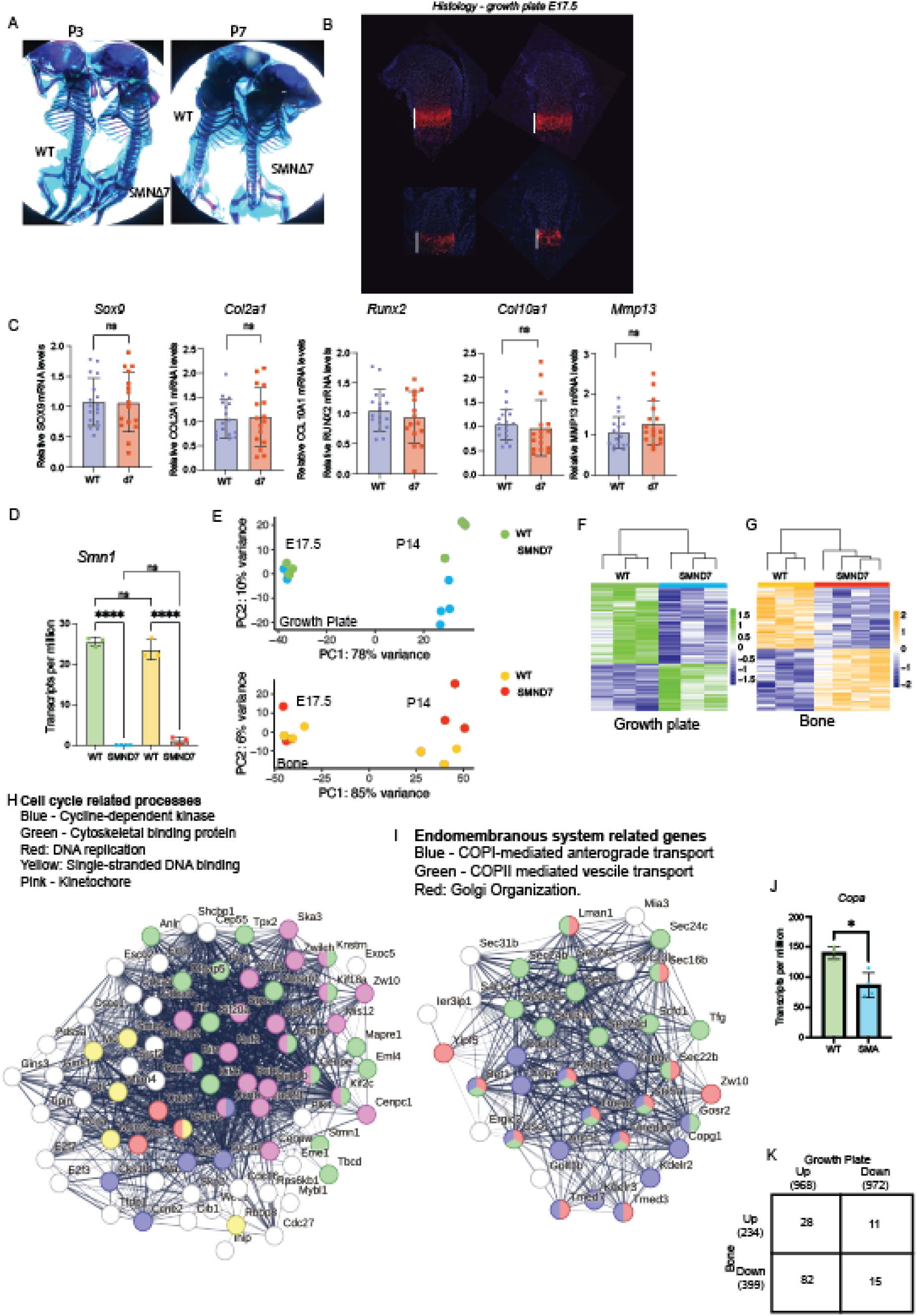
Skeletal Development in the SMNΔ7 mouse model of Type I SMA. **A.** Whole skeletal staining of WT and *SMNΔ7* mice at postnatal day 3 and 7. **B.** Immunofluorescence of the E17.5 growth plate staining for COLX. The length of the COLX positive area is marked with a white line. Not change in the length or the intensity of COLX staining was observed. **C.** Quantitative real-time-PCR on E17.5 growth plates for *Sox9, Col2a1, Runx2, Col10a1,* and *Mmp13.* The mean of both genotypes was tested using a student t-test. **** < 0.0001 **D.** Amount of *Smn1* in P14 growth plate and bone from RNA-sequencing, expressed as transcripts per million. The mean of both genotypes was tested using a student t-test. **** < 0.0001. **E.** PCA analysis of the transcriptomes of E17.5 and P14 growth plates colored by genotype **F.** Heatmaps for growth plate of the differentially expressed genes. Samples are clustered by unsupervised hierarchical clustering. **G.** Heatmaps for bone tissue of the differentially expressed genes. Samples are clustered by unsupervised hierarchical clustering. **H.** STING network map of the downregulated genes in the SMA mouse growth plate associated with cell cycle progression, colored by different cell-cycle terms. **I.** STING network map of the downregulated genes in the SMA mouse growth plate associated with the endomembrane system, colored by which processes, COPI and COPII mediated vesicles and Golgi organization. **J.** Amount of *Copa* in P14 growth plate and bone from RNA-sequencing, expressed as transcripts per million. The mean of both genotypes was tested using a student t-test. * < 0.01. **K.** Comparison between the shared up regulated and downregulated genes at P14 in bone and growth plate.

**Supplemental Figure 3:**
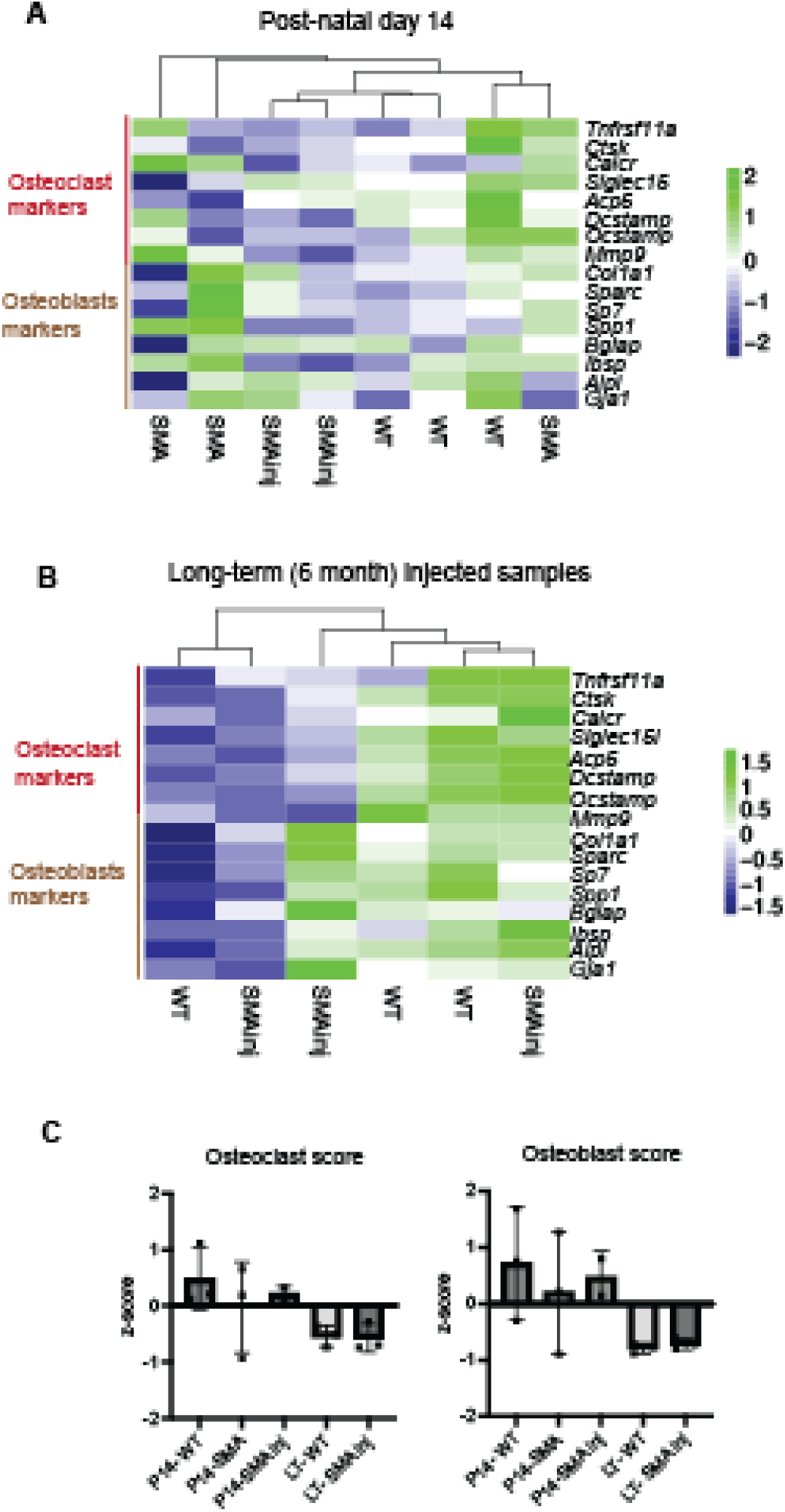
Osteoclast and osteoblast signatures in SMNΔ7 mice. **A.** Heatmap of the osteoclast and osteoblast genes used to make the respective signatures in C in SMA, SMA-injected and WT mice at post-natal day 14, generated from the RNA-sequencing of P14 bone tissue. **B.** Heatmap of the osteoclast and osteoblast genes used to make the respective signatures in C in SMA-injected and WT mice 6 months after injection, generated from the RNA-sequencing of 6-month bone tissue. **C.** Scoring of the osteoclast and osteoblast signatures in each mouse sample represented in A and D. of the SMA, AIS and DMD transcriptomes.

**Supplemental Figure 4:**
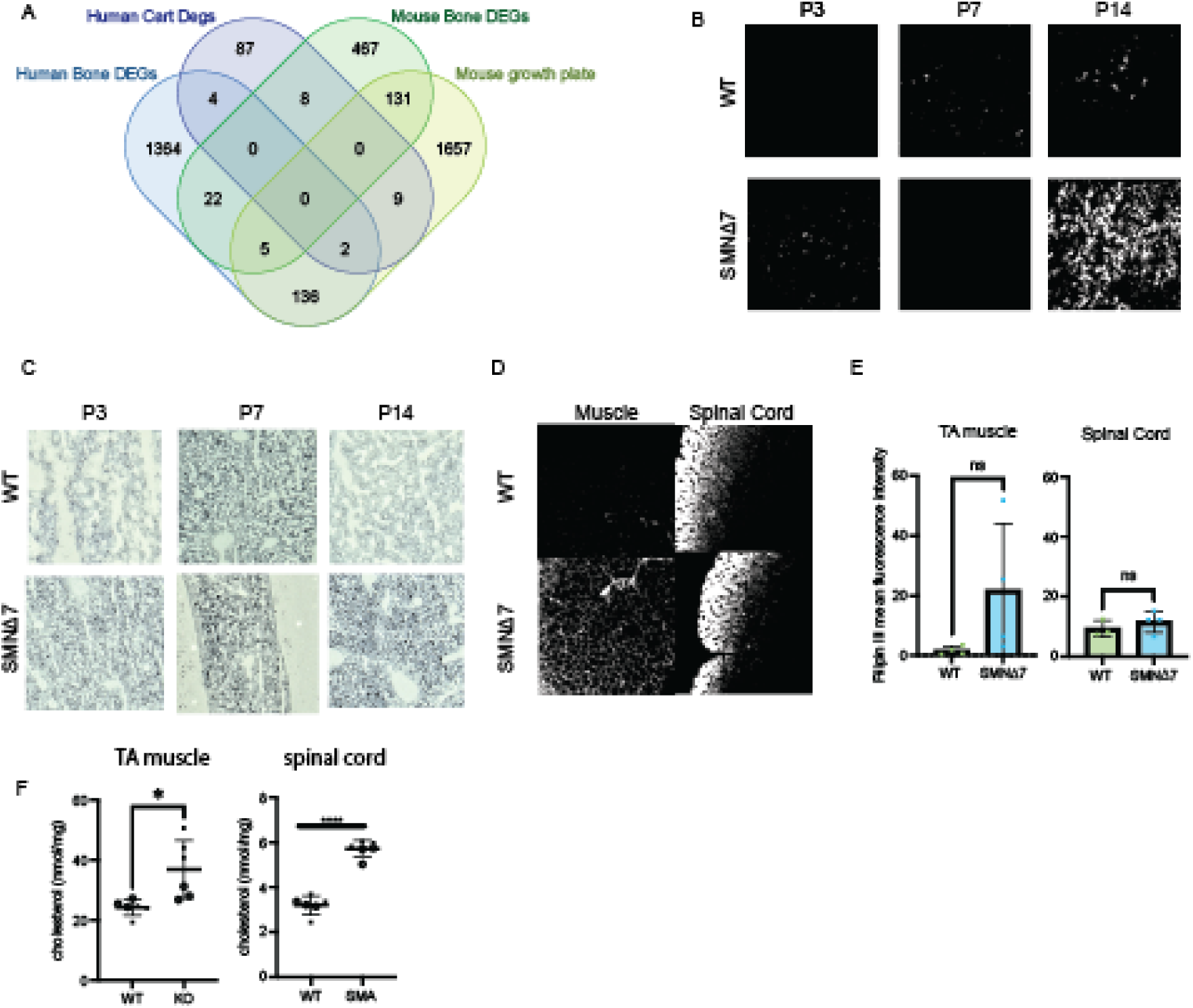
Dynamics of lipid and cholesterol accumulation in SMNΔ7 mouse tissues. **A.** Overlap between the differentially expressed genes in the human and mouse tissues in this study. **B.** Fillipin III staining in the liver at P3, P7 and P14 in WT and SMA animals. **C.** Sudan black staining of the liver at P3, P7 and P14 in WT and SMA animals. **D.** Fillipin III staining at P14 in the muscle (left) and the spinal cord (right). **E.** Quantification of all images taken for the Fillipin staining in D at P14 in the tibialis anterior and spinal cord. Each point represents an animal. Means for each genotype were compared using a student’s t-test. **F.** Quantification of cholesterol via mass-spectrometry in the tibialis anterior and spinal cord at P14 in SMA and WT animals. Means were tested with a student’s t-test. * <0.01, **** <0.0001.

**Supplemental Figure 5:**
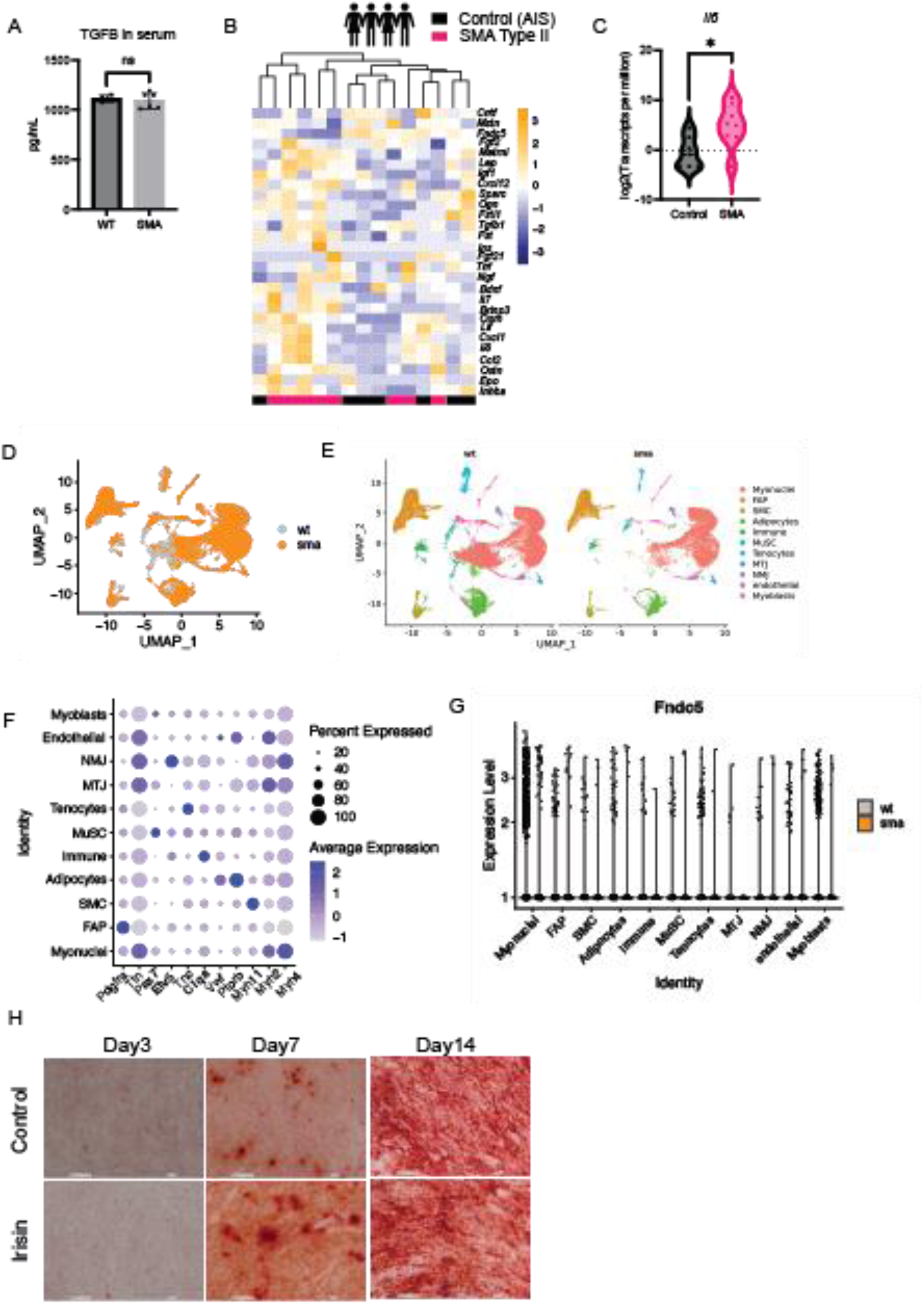
Muscle-derived factors with actions in bone are altered in SMA patients. **A.** ELISA measurement of mouse TGFB1 from the serum of from P14 *SMNΔ7* and WT controls. **B.** Heatmap of myokines in human SMA Type II muscle and AIS controls, remapped from our previous study (*46*). **C.** Violin plot of the transcripts per million (TPMs) of *IL6.* Means are tested only between control and SMA samples using students t-test. n.s – not significant. DMD samples are plot as reference but were not statistically evaluated. **D.** UMAP plot showing the nuclei obtained for the wildtype gastro (grey) from ENCODE and the *SMNΔ7* gastro (orange). Each point represents a single nucleus. **E.** Clusters for each genotype. **F.** Some of the marker genes used to call the cell types in (E). **G.** Expression of *Fndc5* in each cluster of the snRNA-seq dataset from WT and SMA animals, represented as a violin plot. One violin plot is made for the WT and one for the SMA animals. **H.** Representative alizarin red staining images from osteoblast treated with irisin for 3, 7 and 14 days or with vehicle control.

## REFERENCES

1. E. Mercuri, C. J. Sumner, F. Muntoni, B. T. Darras, R. S. Finkel, Spinal muscular atrophy. Nat Rev Dis Primers 8, 1–16 (2022).

2. D. C. Schorling, A. Pechmann, J. Kirschner, Advances in Treatment of Spinal Muscular Atrophy - New Phenotypes, New Challenges, New Implications for Care. J Neuromuscul Dis 7, 1–13 (2020).

3. M. Shababi, C. L. Lorson, S. S. Rudnik-Schöneborn, Spinal muscular atrophy: a motor neuron disorder or a multi-organ disease? Journal of Anatomy 224, 15–28 (2014).

4. C. J. J. Yeo, B. T. Darras, Overturning the Paradigm of Spinal Muscular Atrophy as Just a Motor Neuron Disease. Pediatric Neurology 109, 12–19 (2020).

5. M. D. Mailman, J. W. Heinz, A. C. Papp, P. J. Snyder, M. S. Sedra, B. Wirth, A. H. M. Burghes, T. W. Prior, Molecular analysis of spinal muscular atrophy and modification of the phenotype by SMN2. Genet Med 4, 20–26 (2002).

6. P. Smeriglio, P. Langard, G. Querin, M. G. Biferi, The Identification of Novel Biomarkers Is Required to Improve Adult SMA Patient Stratification, Diagnosis and Treatment. J Pers Med 10, E75 (2020).

7. H. Chaytow, K. M. E. Faller, Y.-T. Huang, T. H. Gillingwater, Spinal muscular atrophy: From approved therapies to future therapeutic targets for personalized medicine. Cell Rep Med 2, 100346 (2021).

8. S. Vai, M. L. Bianchi, I. Moroni, C. Mastella, F. Broggi, L. Morandi, M. T. Arnoldi, C. Bussolino, G. Baranello, Bone and Spinal Muscular Atrophy. Bone 79, 116–120 (2015).

9. H. M. Wasserman, L. N. Hornung, P. J. Stenger, M. M. Rutter, B. L. Wong, I. Rybalsky, J. C. Khoury, H. J. Kalkwarf, Low bone mineral density and fractures are highly prevalent in pediatric patients with spinal muscular atrophy regardless of disease severity. Neuromuscul Disord 27, 331–337 (2017).

10. M. Kinali, L. M. Banks, E. Mercuri, A. Y. Manzur, F. Muntoni, Bone mineral density in a paediatric spinal muscular atrophy population. Neuropediatrics 35, 325–328 (2004).

11. C. Granata, S. Giannini, D. Villa, S. Bonfiglioli Stagni, L. Merlini, Fractures in myopathies. Chir Organi Mov 76, 39–45 (1991).

12. P. Vestergaard, H. Glerup, B. F. Steffensen, L. Rejnmark, J. Rahbek, L. Moseklide, Fracture risk in patients with muscular dystrophy and spinal muscular atrophy. J Rehabil Med 33, 150–155 (2001).

13. A. Fujak, C. Kopschina, R. Forst, F. Gras, L. A. Mueller, J. Forst, Fractures in proximal spinal muscular atrophy. Arch Orthop Trauma Surg 130, 775–780 (2010).

14. J. Y.-L. Tung, T.-K. Chow, M. Wai, J. Lo, S. H. S. Chan, Bone Health Status of Children with Spinal Muscular Atrophy. Journal of Bone Metabolism 30, 319 (2023).

15. A.-K. Kroksmark, L. Alberg, M. Tulinius, P. Magnusson, A.-C. Söderpalm, Low bone mineral density and reduced bone-specific alkaline phosphatase in 5q spinal muscular atrophy type 2 and type 3: A 2-year prospective study of bone health. Acta Paediatrica 112, 2589–2600 (2023).

16. A. L. Xu, T. O. Crawford, P. D. Sponseller, Hip Pain in Patients With Spinal Muscular Atrophy: Prevalence, Intensity, Interference, and Factors Associated With Moderate to Severe Pain. J Pediatr Orthop 42, 273–279 (2022).

17. R. B. Hanna, N. Nahm, M. A. Bent, S. Sund, K. Patterson, M. K. Schroth, M. A. Halanski, Hip Pain in Nonambulatory Children with Type-I or II Spinal Muscular Atrophy. JB JS Open Access 7, e22.00011 (2022).

18. S. M. Sporer, B. G. Smith, Hip Dislocation in Patients With Spinal Muscular Atrophy. Journal of Pediatric Orthopaedics 23, 10 (2003).

19. C. Granata, L. Merlini, E. Magni, M. L. Marini, S. B. Stagni, Spinal muscular atrophy: natural history and orthopaedic treatment of scoliosis. Spine (Phila Pa 1976) 14, 760–762 (1989).

20. E. Rodillo, M. L. Marini, J. Z. Heckmatt, V. Dubowitz, Scoliosis in Spinal Muscular Atrophy: Review of 63 Cases. J Child Neurol 4, 118–123 (1989).

21. D. P. Phillips, D. P. Roye, J. P. Farcy, A. Leet, Y. A. Shelton, Surgical treatment of scoliosis in a spinal muscular atrophy population. Spine (Phila Pa 1976) 15, 942–945 (1990).

22. C. A. Wijngaarde, R. C. Brink, F. A. S. de Kort, M. Stam, L. A. M. Otto, F.-L. Asselman, B. Bartels, R. P. A. van Eijk, J. Sombroek, I. Cuppen, M. Verhoef, L. H. van den Berg, R. I. Wadman, R. M. Castelein, W.-L. van der Pol, Natural course of scoliosis and lifetime risk of scoliosis surgery in spinal muscular atrophy. Neurology 93, e149–e158 (2019).

23. A. Fujak, W. Raab, A. Schuh, S. Richter, R. Forst, J. Forst, Natural course of scoliosis in proximal spinal muscular atrophy type II and IIIa: descriptive clinical study with retrospective data collection of 126 patients. BMC Musculoskelet Disord 14, 283 (2013).

24. K. E. Chad, H. A. McKay, G. A. Zello, D. A. Bailey, R. A. Faulkner, R. E. Snyder, Body composition in nutritionally adequate ambulatory and non-ambulatory children with cerebral palsy and a healthy reference group. Dev Med Child Neurol 42, 334–339 (2000).

25. G. Baranello, S. Vai, F. Broggi, R. Masson, M. T. Arnoldi, R. Zanin, C. Mastella, M. L. Bianchi, Evolution of bone mineral density, bone metabolism and fragility fractures in Spinal Muscular Atrophy (SMA) types 2 and 3. Neuromuscular Disorders 29, 525–532 (2019).

26. N. J. Crabtree, H. Roper, N. J. Shaw, Cessation of ambulation results in a dramatic loss of trabecular bone density in boys with Duchenne muscular dystrophy (DMD). Bone 154, 116248 (2022).

27. I. A. Khatri, U. S. Chaudhry, M. G. Seikaly, R. H. Browne, S. T. Iannaccone, Low Bone Mineral Density in Spinal Muscular Atrophy. Journal of Clinical Neuromuscular Disease 10, 11–17 (2008).

28. D. T. Kobayashi, J. Shi, L. Stephen, K. L. Ballard, R. Dewey, J. Mapes, B. Chung, K. McCarthy, K. J. Swoboda, T. O. Crawford, R. Li, T. Plasterer, C. Joyce, the B. for S. M. A. S. Group, W. K. Chung, P. Kaufmann, B. T. Darras, R. S. Finkel, D. M. Sproule, W. B. Martens, M. P. McDermott, D. C. D. Vivo, the P. N. C. R. Network, M. G. Walker, K. S. Chen, SMA-MAP: A Plasma Protein Panel for Spinal Muscular Atrophy. PLOS ONE 8, e60113 (2013).

29. N. E. Yasar, G. Ozdemir, E. Uzun Ata, M. O. Ayvali, N. Ata, M. Ulgu, E. Dumlupınar, S. Birinci, I. Bingol, S. Bekmez, Nusinersen therapy changed the natural course of spinal muscular atrophy type 1: What about spine and hip? Journal of Children’s Orthopaedics 18, 322–330 (2024).

30. F. Al Amrani, R. Amin, J. Chiang, L. Xiao, J. Boyd, E. Law, E. Nigro, L. Weinstock, A. Stosic, H. D. Gonorazky, Scoliosis in Spinal Muscular Atrophy Type 1 in the Nusinersen Era. Neurol Clin Pract 12, 279–287 (2022).

31. V. Soini, G. Schreiber, B. Wilken, A. K. Hell, Early Development of Spinal Deformities in Children Severely Affected with Spinal Muscular Atrophy after Gene Therapy with Onasemnogene Abeparvovec—Preliminary Results. Children (Basel) 10, 998 (2023).

32. S.-H. Hann, S.-Y. Kim, Y.-L. Kim, Y.-W. Jo, J.-S. Kang, H. Park, S.-Y. Choi, Y.-Y. Kong, Depletion of SMN Protein in Mesenchymal Progenitors Impairs the Development of Bone and Neuromuscular Junction in Spinal Muscular Atrophy. eLife 12 (2023).

33. N. Hensel, H. Brickwedde, K. Tsaknakis, A. Grages, L. Braunschweig, K. A. Lüders, H. M. Lorenz, S. Lippross, L. M. Walter, F. Tavassol, S. Lienenklaus, C. Neunaber, P. Claus, A. K. Hell, Altered bone development with impaired cartilage formation precedes neuromuscular symptoms in spinal muscular atrophy. Human Molecular Genetics 29, 2662–2673 (2020).

34. S. Shanmugarajan, E. Tsuruga, K. J. Swoboda, B. L. Maria, W. L. Ries, S. V. Reddy, Bone loss in survival motor neuron (Smn−/−SMN2) genetic mouse model of spinal muscular atrophy. J Pathol 219, 52–60 (2009).

35. S. Shanmugarajan, K. J. Swoboda, S. T. Iannaccone, W. L. Ries, B. L. Maria, S. V. Reddy, Congenital bone fractures in spinal muscular atrophy: functional role for SMN protein in bone remodeling. J Child Neurol 22, 967–973 (2007).

36. N. Kurihara, C. Menaa, H. Maeda, D. J. Haile, S. V. Reddy, Osteoclast-stimulating factor interacts with the spinal muscular atrophy gene product to stimulate osteoclast formation. J Biol Chem 276, 41035–41039 (2001).

37. SMA Europe | Priority Setting Project | Shaping SMA Initiatives. https://www.sma-europe.eu/our-priority-setting-project.

38. L. Miladi, M. Gaume, N. Khouri, M. Johnson, V. Topouchian, C. Glorion, Minimally Invasive Surgery for Neuromuscular Scoliosis: Results and Complications in a Series of One Hundred Patients. Spine 43, E968 (2018).

39. M. Gaume, E. Saudeau, M. Gomez-Garcia de la Banda, V. Azzi-Salameh, B. Mbieleu, D. Verollet, A. Benezit, J. Bergounioux, A. Essid, I. Doehring, I. Dabaj, I. Desguerre, C. Barnerias, V. Topouchian, C. Glorion, S. Quijano-Roy, L. Miladi, Minimally Invasive Fusionless Surgery for Scoliosis in Spinal Muscular Atrophy: Long-term Follow-up Results in a Series of 59 Patients. J Pediatr Orthop 41, 549–558 (2021).

40. M. Gaumé, E. Saghbiny, L. Richard, C. Thouement, R. Vialle, L. Miladi, Pelvic Fixation Technique Using the Ilio-Sacral Screw for 173 Neuromuscular Scoliosis Patients. Children (Basel) 11, 199 (2024).

41. T. Langlais, A. Bouy, G. Eloy, N. Mainard, W. Skalli, C. Vergari, R. Vialle, Sagittal plane assessment of manual concave rod bending for posterior correction in adolescents with idiopathic thoracic scoliosis (Lenke 1 and 3). Orthop Traumatol Surg Res 109, 103654 (2023).

42. J. L. Buckner, S. A. Bowden, J. D. Mahan, Optimizing Bone Health in Duchenne Muscular Dystrophy. Int J Endocrinol 2015, 928385 (2015).

43. S. Bandyopadhyay, M. P. Duffy, K. J. Ahn, J. H. Sussman, M. Pang, D. Smith, G. Duncan, I. Zhang, J. Huang, Y. Lin, B. Xiong, T. Imtiaz, C.-H. Chen, A. Thadi, C. Chen, J. Xu, M. Reichart, Z. Martinez, C. Diorio, C. Chen, V. Pillai, O. Snaith, D. Oldridge, S. Bhattacharyya, I. Maillard, M. Carroll, C. Nelson, L. Qin, K. Tan, Mapping the cellular biogeography of human bone marrow niches using single-cell transcriptomics and proteomic imaging. Cell 187, 3120–3140.e29 (2024).

44. Novartis/granulator, Novartis (2024); https://github.com/Novartis/granulator.

45. G. J. Hunt, S. Freytag, M. Bahlo, J. A. Gagnon-Bartsch, dtangle: accurate and robust cell type deconvolution. Bioinformatics 35, 2093–2099 (2019).

46. F. C. Grandi, S. Astord, S. Pezet, E. Gidaja, S. Mazzucchi, M. Chapart, S. Vasseur, K. Mamchaoui, P. Smeriglio, SMA Type II Skeletal Muscle Treated with Nusinersen shows SMN Restoration but Mitochondrial Deficiency. bioRxiv [Preprint] (2024). 10.1101/2024.02.29.582680.

47. F. Gabanella, C. Pisani, A. Borreca, S. Farioli-Vecchioli, M. T. Ciotti, T. Ingegnere, A. Onori, M. Ammassari-Teule, N. Corbi, N. Canu, L. Monaco, C. Passananti, M. G. Di Certo, SMN affects membrane remodelling and anchoring of the protein synthesis machinery. Journal of Cell Science 129, 804–816 (2016).

48. S. K. Custer, J. W. Astroski, H. X. Li, E. J. Androphy, Interaction between alpha-COP and SMN ameliorates disease phenotype in a mouse model of spinal muscular atrophy. Biochem Biophys Res Commun 514, 530–537 (2019).

49. H. Li, S. K. Custer, T. Gilson, L. T. Hao, C. E. Beattie, E. J. Androphy, α-COP binding to the survival motor neuron protein SMN is required for neuronal process outgrowth. Hum Mol Genet 24, 7295–7307 (2015).

50. R. D. Labrom, Growth and Maturation of the Spine from Birth to Adolescence. JBJS 89, 3 (2007).

51. I. A. F. Stokes, H. Spence, D. D. Aronsson, N. Kilmer, Mechanical Modulation of Vertebral Body Growth: Implications for Scoliosis Progression. Spine 21, 1162 (1996).

52. S. Monteagudo, F. M. F. Cornelis, C. Aznar-Lopez, P. Yibmantasiri, L.-A. Guns, P. Carmeliet, F. Cailotto, R. J. Lories, DOT1L safeguards cartilage homeostasis and protects against osteoarthritis. Nat Commun 8, 15889 (2017).

53. M. E. R. Butchbach, J. D. Edwards, A. H. M. Burghes, Abnormal motor phenotype in the SMNΔ7 mouse model of spinal muscular atrophy. Neurobiol Dis 27, 207–219 (2007).

54. T. T. Le, L. T. Pham, M. E. R. Butchbach, H. L. Zhang, U. R. Monani, D. D. Coovert, T. O. Gavrilina, L. Xing, G. J. Bassell, A. H. M. Burghes, SMNΔ7, the major product of the centromeric survival motor neuron (SMN2) gene, extends survival in mice with spinal muscular atrophy and associates with full-length SMN. Human Molecular Genetics 14, 845–857 (2005).

55. E. C. Weir, W. M. Philbrick, M. Amling, L. A. Neff, R. Baron, A. E. Broadus, Targeted overexpression of parathyroid hormone-related peptide in chondrocytes causes chondrodysplasia and delayed endochondral bone formation. Proc Natl Acad Sci U S A 93, 10240–10245 (1996).

56. L. Wang, Y. Y. Shao, R. T. Ballock, Peroxisome Proliferator-Activated Receptor-gamma Promotes Adipogenic Changes in Growth Plate Chondrocytes In Vitro. PPAR Res 2006, 67297 (2006).

57. E. Dominguez, T. Marais, N. Chatauret, S. Benkhelifa-Ziyyat, S. Duque, P. Ravassard, R. Carcenac, S. Astord, A. Pereira de Moura, T. Voit, M. Barkats, Intravenous scAAV9 delivery of a codon-optimized SMN1 sequence rescues SMA mice. Hum Mol Genet 20, 681–693 (2011).

58. A. Besse, S. Astord, T. Marais, M. Roda, B. Giroux, F.-X. Lejeune, F. Relaix, P. Smeriglio, M. Barkats, M. G. Biferi, AAV9-Mediated Expression of SMN Restricted to Neurons Does Not Rescue the Spinal Muscular Atrophy Phenotype in Mice. Mol Ther 28, 1887–1901 (2020).

59. D. M. Ramos, C. d’Ydewalle, V. Gabbeta, A. Dakka, S. K. Klein, D. A. Norris, J. Matson, S. J. Taylor, P. G. Zaworski, T. W. Prior, P. J. Snyder, D. Valdivia, C. L. Hatem, I. Waters, N. Gupte, K. J. Swoboda, F. Rigo, C. F. Bennett, N. Naryshkin, S. Paushkin, T. O. Crawford, C. J. Sumner, Age-dependent SMN expression in disease-relevant tissue and implications for SMA treatment. J Clin Invest 129, 4817–4831 (2019).

60. S. Bolamperti, I. Villa, A. Rubinacci, Bone remodeling: an operational process ensuring survival and bone mechanical competence. Bone Res 10, 1–19 (2022).

61. M. M. Weivoda, C. K. Chew, D. G. Monroe, J. N. Farr, E. J. Atkinson, J. R. Geske, B. Eckhardt, B. Thicke, M. Ruan, A. J. Tweed, L. K. McCready, R. A. Rizza, A. Matveyenko, M. Kassem, T. L. Andersen, A. Vella, M. T. Drake, B. L. Clarke, M. J. Oursler, S. Khosla, Identification of osteoclast-osteoblast coupling factors in humans reveals links between bone and energy metabolism. Nat Commun 11, 87 (2020).

62. K. Fuller, K. E. Bayley, T. J. Chambers, Activin A is an essential cofactor for osteoclast induction. Biochem Biophys Res Commun 268, 2–7 (2000).

63. T. Kajita, W. Ariyoshi, T. Okinaga, S. Mitsugi, K. Tominaga, T. Nishihara, Mechanisms involved in enhancement of osteoclast formation by activin-A. Journal of Cellular Biochemistry 119, 6974–6985 (2018).

64. R. S. Pearsall, E. Canalis, M. Cornwall-Brady, K. W. Underwood, B. Haigis, J. Ucran, R. Kumar, E. Pobre, A. Grinberg, E. D. Werner, V. Glatt, L. Stadmeyer, D. Smith, J. Seehra, M. L. Bouxsein, A soluble activin Type IIA receptor induces bone formation and improves skeletal integrity. Proceedings of the National Academy of Sciences 105, 7082–7087 (2008).

65. R. Sakai, Y. Eto, M. Ohtsuka, M. Hirafuji, H. Shinoda, Activin Enhances Osteoclast-like Cell Formation *in Vitro*. Biochemical and Biophysical Research Communications 195, 39–46 (1993).

66. S. Vallet, S. Mukherjee, N. Vaghela, T. Hideshima, M. Fulciniti, S. Pozzi, L. Santo, D. Cirstea, K. Patel, A. R. Sohani, A. Guimaraes, W. Xie, D. Chauhan, J. A. Schoonmaker, E. Attar, M. Churchill, E. Weller, N. Munshi, J. S. Seehra, R. Weissleder, K. C. Anderson, D. T. Scadden, N. Raje, Activin A promotes multiple myeloma-induced osteolysis and is a promising target for myeloma bone disease. Proceedings of the National Academy of Sciences 107, 5124–5129 (2010).

67. Y. Cao, I. D. C. Jansen, S. Sprangers, J. Stap, P. J. M. Leenen, V. Everts, T. J. de Vries, IL-1β differently stimulates proliferation and multinucleation of distinct mouse bone marrow osteoclast precursor subsets. J Leukoc Biol 100, 513–523 (2016).

68. A. Levescot, M. H. Chang, J. Schnell, N. Nelson-Maney, J. Yan, M. Martínez-Bonet, R. Grieshaber-Bouyer, P. Y. Lee, K. Wei, R. B. Blaustein, A. Morris, A. Wactor, Y. Iwakura, J. A. Lederer, D. A. Rao, J. F. Charles, P. A. Nigrovic, IL-1β-driven osteoclastogenic Tregs accelerate bone erosion in arthritis. J Clin Invest 131, e141008 (2021).

69. Z. Chen, K. Buki, J. Vääräniemi, G. Gu, H. K. Väänänen, The critical role of IL-34 in osteoclastogenesis. PLoS One 6, e18689 (2011).

70. X. Wu, C. Yang, X. Chen, Z. Shan, X. Wu, Interferon Regulatory Factor 4 (IRF4) Plays a Key Role in Osteoblast Differentiation of Postmenopausal Osteoporosis. Front Biosci (Landmark Ed) 29, 115 (2024).

71. Y. Nakashima, T. Haneji, Stimulation of osteoclast formation by RANKL requires interferon regulatory factor-4 and is inhibited by simvastatin in a mouse model of bone loss. PLoS One 8, e72033 (2013).

72. E. Y. Chen, C. M. Tan, Y. Kou, Q. Duan, Z. Wang, G. V. Meirelles, N. R. Clark, A. Ma’ayan, Enrichr: interactive and collaborative HTML5 gene list enrichment analysis tool. BMC Bioinformatics 14, 128 (2013).

73. N. S. Alekos, M. C. Moorer, R. C. Riddle, Dual Effects of Lipid Metabolism on Osteoblast Function. Front. Endocrinol. 11 (2020).

74. E. Rendina-Ruedy, C. J. Rosen, Lipids in the Bone Marrow: An Evolving Perspective. Cell Metabolism 31, 219–231 (2020).

75. B. Wang, H. Wang, Y. Li, L. Song, Lipid metabolism within the bone micro-environment is closely associated with bone metabolism in physiological and pathophysiological stages. Lipids Health Dis 21, 5 (2022).

76. P. E. Pardee, W. Dunn, J. B. Schwimmer, Nonalcoholic Fatty Liver Disease is Associated with Low Bone Mineral Density in Obese Children. Aliment Pharmacol Ther 35, 248–254 (2012).

77. T. Sato, I. Morita, S. Murota, Involvement of cholesterol in osteoclast-like cell formation via cellular fusion. Bone 23, 135–140 (1998).

78. E. Luegmayr, H. Glantschnig, G. A. Wesolowski, M. A. Gentile, J. E. Fisher, G. A. Rodan, A. A. Reszka, Osteoclast formation, survival and morphology are highly dependent on exogenous cholesterol/lipoproteins. Cell Death Differ 11, S108–S118 (2004).

79. Y. Feng, Y. Cui, J. Jin, S. Huang, J. Wei, M. Yao, D. Zhou, S. Mao, The Alterations of Gut Microbiome and Lipid Metabolism in Patients with Spinal Muscular Atrophy. Neurol Ther 12, 961–976 (2023).

80. D. M.-K. Leow, Y. K. Ng, L. C. Wang, H. W. L. Koh, T. Zhao, Z. J. Khong, T. Tabaglio, G. Narayanan, R. M. Giadone, R. M. Sobota, S.-Y. Ng, A. K. K. Teo, S. H. Parson, L. L. Rubin, W.-Y. Ong, B. T. Darras, C. J. J. Yeo, Hepatocyte-intrinsic SMN deficiency drives metabolic dysfunction and liver steatosis in spinal muscular atrophy. J Clin Invest 134 (2024).

81. E. R. Sutton, A. Beauvais, R. Yaworski, Y. D. Repentigny, A. Reilly, M. M. A. de Almeida, M.-O. Deguise, K. L. Poulin, R. J. Parks, B. L. Schneider, R. Kothary, Liver SMN restoration rescues the Smn2B/- mouse model of spinal muscular atrophy. eBioMedicine 110 (2024).

82. M. Deguise, G. Baranello, C. Mastella, A. Beauvais, J. Michaud, A. Leone, R. De Amicis, A. Battezzati, C. Dunham, K. Selby, J. Warman Chardon, H. J. McMillan, Y. Huang, N. L. Courtney, A. J. Mole, S. Kubinski, P. Claus, L. M. Murray, M. Bowerman, T. H. Gillingwater, S. Bertoli, S. H. Parson, R. Kothary, Abnormal fatty acid metabolism is a core component of spinal muscular atrophy. Ann Clin Transl Neurol 6, 1519–1532 (2019).

83. M.-O. Deguise, L. Chehade, A. Tierney, A. Beauvais, R. Kothary, Low fat diets increase survival of a mouse model of spinal muscular atrophy. Annals of Clinical and Translational Neurology 6, 2340–2346 (2019).

84. M. Bowerman, K. J. Swoboda, J.-P. Michalski, G.-S. Wang, C. Reeks, A. Beauvais, K. Murphy, J. Woulfe, R. A. Screaton, F. W. Scott, R. Kothary, Glucose metabolism and pancreatic defects in spinal muscular atrophy. Annals of Neurology 72, 256–268 (2012).

85. A. Reilly, A. Beauvais, M. Al-Aarg, R. Yaworski, E. R. Sutton, S. Thebault, R. Kothary, Peripheral defects precede neuromuscular pathology in the Smn2B/− mouse model of spinal muscular atrophy. Journal of Neuromuscular Diseases 11, 1200–1210 (2024).

86. M. M. Syed-Abdul, Lipid Metabolism in Metabolic-Associated Steatotic Liver Disease (MASLD). Metabolites 14, 12 (2024).

87. S. Rath, R. Sharma, R. Gupta, T. Ast, C. Chan, T. J. Durham, R. P. Goodman, Z. Grabarek, M. E. Haas, W. H. W. Hung, P. R. Joshi, A. A. Jourdain, S. H. Kim, A. V. Kotrys, S. S. Lam, J. G. McCoy, J. D. Meisel, M. Miranda, A. Panda, A. Patgiri, R. Rogers, S. Sadre, H. Shah, O. S. Skinner, T.-L. To, M. A. Walker, H. Wang, P. S. Ward, J. Wengrod, C.-C. Yuan, S. E. Calvo, V. K. Mootha, MitoCarta3.0: an updated mitochondrial proteome now with sub-organelle localization and pathway annotations. Nucleic Acids Res 49, D1541–D1547 (2021).

88. P. F. Dobson, E. P. Dennis, D. Hipps, A. Reeve, A. Laude, C. Bradshaw, C. Stamp, A. Smith, D. J. Deehan, D. M. Turnbull, L. C. Greaves, Mitochondrial dysfunction impairs osteogenesis, increases osteoclast activity, and accelerates age related bone loss. Sci Rep 10, 11643 (2020).

89. T. R. Koves, J. R. Ussher, R. C. Noland, D. Slentz, M. Mosedale, O. Ilkayeva, J. Bain, R. Stevens, J. R. B. Dyck, C. B. Newgard, G. D. Lopaschuk, D. M. Muoio, Mitochondrial Overload and Incomplete Fatty Acid Oxidation Contribute to Skeletal Muscle Insulin Resistance. Cell Metabolism 7, 45–56 (2008).

90. R. Sheng, M. Cao, M. Song, M. Wang, Y. Zhang, L. Shi, T. Xie, Y. Li, J. Wang, Y. Rui, Muscle-bone crosstalk via endocrine signals and potential targets for osteosarcopenia-related fracture. Journal of Orthopaedic Translation 43, 36–46 (2023).

91. Y. Mao, Z. Jin, J. Yang, D. Xu, L. Zhao, A. Kiram, Y. Yin, D. Zhou, Z. Sun, L. Xiao, Z. Zhou, L. Yang, T. Fu, Z. Xu, Y. Jia, X. Chen, F.-N. Niu, X. Li, Z. Zhu, Z. Gan, Muscle-bone cross-talk through the FNIP1-TFEB-IGF2 axis is associated with bone metabolism in human and mouse. Sci Transl Med 16, eadk9811 (2024).

92. M. Shao, Q. Wang, Q. Lv, Y. Zhang, G. Gao, S. Lu, Advances in the research on myokine-driven regulation of bone metabolism. Heliyon 10, e22547 (2024).

93. H. Kaji, Effects of myokines on bone. Bonekey Rep 5, 826 (2016).

94. A. Brener, L. Sagi, A. Shtamler, S. Levy, A. Fattal-Valevski, Y. Lebenthal, Insulin-like growth factor-1 status is associated with insulin resistance in young patients with spinal muscular atrophy. Neuromuscul Disord 30, 888–896 (2020).

95. A. Y. Kaymaz, S. K. Bal, G. Bora, B. Talim, A. Ozon, A. Alikasifoglu, H. Topaloglu, H. E. Yurter, Alterations in insulin-like growth factor system in spinal muscular atrophy. Muscle Nerve 66, 631–638 (2022).

96. L.-K. Tsai, Y.-C. Chen, W.-C. Cheng, C.-H. Ting, J. C. Dodge, W.-L. Hwu, S. H. Cheng, M. A. Passini, IGF-1 delivery to CNS attenuates motor neuron cell death but does not improve motor function in type III SMA mice. Neurobiology of Disease 45, 272–279 (2012).

97. L.-K. Tsai, C.-L. Chen, C.-H. Ting, S. Lin-Chao, W.-L. Hwu, J. C. Dodge, M. A. Passini, S. H. Cheng, Systemic administration of a recombinant AAV1 vector encoding IGF-1 improves disease manifestations in SMA mice. Mol Ther 22, 1450–1459 (2014).

98. M. Karlsson, C. Zhang, L. Méar, W. Zhong, A. Digre, B. Katona, E. Sjöstedt, L. Butler, J. Odeberg, P. Dusart, F. Edfors, P. Oksvold, K. von Feilitzen, M. Zwahlen, M. Arif, O. Altay, X. Li, M. Ozcan, A. Mardinoglu, L. Fagerberg, J. Mulder, Y. Luo, F. Ponten, M. Uhlén, C. Lindskog, A single–cell type transcriptomics map of human tissues. Science Advances 7, eabh2169 (2021).

99. FNDC5 protein expression summary - The Human Protein Atlas. https://www.proteinatlas.org/ENSG00000160097-FNDC5.

100. ENCBS759QMD – ENCODE. https://www.encodeproject.org/biosamples/ENCBS759QMD/.

101. M. A, T. E. Wales, H. Zhou, S.-V. Draga-Coletă, C. Gorgulla, K. A. Blackmore, M. J. Mittenbühler, C. R. Kim, D. Bogoslavski, Q. Zhang, Z.-F. Wang, M. P. Jedrychowski, H.-S. Seo, K. Song, A. Z. Xu, L. Sebastian, S. P. Gygi, H. Arthanari, S. Dhe-Paganon, P. R. Griffin, J. R. Engen, B. M. Spiegelman, Irisin acts through its integrin receptor in a two-step process involving extracellular Hsp90α. Molecular Cell 83, 1903–1920.e12 (2023).

102. H. Kim, C. D. Wrann, M. Jedrychowski, S. Vidoni, Y. Kitase, K. Nagano, C. Zhou, J. Chou, V.-J. A. Parkman, S. J. Novick, T. S. Strutzenberg, B. D. Pascal, P. T. Le, D. J. Brooks, A. M. Roche, K. K. Gerber, L. Mattheis, W. Chen, H. Tu, M. L. Bouxsein, P. R. Griffin, R. Baron, C. J. Rosen, L. F. Bonewald, B. M. Spiegelman, Irisin Mediates Effects on Bone and Fat via αV Integrin Receptors. Cell 175, 1756–1768.e17 (2018).

103. G. Colaianni, C. Cuscito, T. Mongelli, P. Pignataro, C. Buccoliero, P. Liu, P. Lu, L. Sartini, M. Di Comite, G. Mori, A. Di Benedetto, G. Brunetti, T. Yuen, L. Sun, J. E. Reseland, S. Colucci, M. I. New, M. Zaidi, S. Cinti, M. Grano, The myokine irisin increases cortical bone mass. Proceedings of the National Academy of Sciences 112, 12157–12162 (2015).

104. E. Solsona-Vilarrasa, R. Fucho, S. Torres, S. Nuñez, N. Nuño-Lámbarri, C. Enrich, C. García-Ruiz, J. C. Fernández-Checa, Cholesterol enrichment in liver mitochondria impairs oxidative phosphorylation and disrupts the assembly of respiratory supercomplexes. Redox Biol 24, 101214 (2019).

105. F. Amor, A. Vu Hong, G. Corre, M. Sanson, L. Suel, S. Blaie, L. Servais, T. Voit, I. Richard, D. Israeli, Cholesterol metabolism is a potential therapeutic target in Duchenne muscular dystrophy. Journal of Cachexia, Sarcopenia and Muscle 12, 677–693 (2021).

106. H. J. Lim, H. Yoon, J. Kim, K. Han, Y. So, M. Park, K.-B. Park, M.-J. Lee, Comparison of elasticity changes in the paraspinal muscles of adolescent patients with scoliosis treated with surgery and bracing. Sci Rep 14, 5623 (2024).

107. H. Ye, Y. Xu, R. Mi, Y. Liu, Y. Lyu, S. Wu, G. Wu, Evaluation of Paravertebral Muscle Structure Asymmetry in Idiopathic Scoliosis Using Imaging Techniques. World Neurosurgery 191, e547–e555 (2024).

108. A. F. Mannion, M. Meier, D. Grob, M. Müntener, Paraspinal muscle fibre type alterations associated with scoliosis: an old problem revisited with new evidence. Eur Spine J 7, 289–293 (1998).

109. E. Bal, S. Batin, Comparison of morphometric measurements of lumbar muscles on the convex and concave sides of curvature in idiopathic scoliosis. Medicine (Baltimore) 102, e35667 (2023).

110. N. A. Sims, Influences of the IL-6 cytokine family on bone structure and function. Cytokine 146, 155655 (2021).

111. F. Blanchard, L. Duplomb, M. Baud’huin, B. Brounais, The dual role of IL-6-type cytokines on bone remodeling and bone tumors. Cytokine Growth Factor Rev 20, 19–28 (2009).

112. D. Rigueur, K. M. Lyons, Whole-mount skeletal staining. Methods Mol Biol 1130, 113–121 (2014).

113. F. C. Grandi, H. Modi, L. Kampman, M. R. Corces, Chromatin accessibility profiling by ATAC-seq. Nat Protoc 17, 1518–1552 (2022).

114. M. D. Young, S. Behjati, SoupX removes ambient RNA contamination from droplet-based single-cell RNA sequencing data. GigaScience 9, giaa151 (2020).

115. C. S. McGinnis, L. M. Murrow, Z. J. Gartner, DoubletFinder: Doublet Detection in Single-Cell RNA Sequencing Data Using Artificial Nearest Neighbors. cels 8, 329–337.e4 (2019).

116. C. Chevalier, M. Çolakoğlu, J. Brun, C. Thouverey, N. Bonnet, S. Ferrari, M. Trajkovski, Primary mouse osteoblast and osteoclast culturing and analysis. STAR Protoc 2, 100452 (2021).

117. H. Hojo, T. Saito, X. He, Q. Guo, S. Onodera, T. Azuma, M. Koebis, K. Nakao, A. Aiba, M. Seki, Y. Suzuki, H. Okada, S. Tanaka, U. Chung, A. P. McMahon, S. Ohba, Runx2 regulates chromatin accessibility to direct the osteoblast program at neonatal stages. Cell Reports 40, 111315 (2022).

